# Dynamically Linking Influenza Virus Infection Kinetics, Lung Injury, Inflammation, and Disease Severity

**DOI:** 10.1101/555276

**Authors:** Margaret A. Myers, Amanda P. Smith, Lindey C. Lane, David J. Moquin, Rosemary Aogo, Stacie Woolard, Paul G. Thomas, Peter Vogel, Amber M. Smith

## Abstract

Influenza viruses cause a significant amount of morbidity and mortality. Understanding host immune control efficacy and how different factors influence lung injury and disease severity are critical. Here, we established dynamical connections between viral loads, infected cells, CD8^+^ T cell-mediated clearance, lung injury, inflammation, and disease severity using an integrative model-experiment exchange. The model was validated through CD8 depletion and whole lung histomorphometry, which showed that the infected area matched the model-predicted infected cell dynamics and that the resolved area paralleled the relative CD8 dynamics. Inflammation could further be predicted by the infected cell dynamics, and additional analyses revealed nonlinear relations between lung injury, inflammation, and disease severity. These links between important pathogen kinetics and host pathology enhance our ability to forecast disease progression, potential complications, and therapeutic efficacy.

## Introduction

Over 15 million respiratory infections and 200,000 hospitalizations result from influenza A viruses (IAVs) each year [1–4]. The incidence and severity of IAV infections increases when new strains emerge and/or when there is a lack of prior immunity. A robust immune response is crucial for resolving viral infections, but immune-mediated pathology can exacerbate disease [5–9]. High viral loads also play a role in disease progression [10], but these do not always correlate with the strength of the host response or with disease severity [11–14]. An understanding of how viral loads, host immune responses, and disease progression are related is critical to identify disease-specific markers that may help predict hospitalization or other complications.

During IAV infection, viral loads increase rapidly for the first 1-2 days of infection before reaching a peak (e.g., as in [13,15–19]). In naïve hosts, viral loads then begin to decline, first slowly (< 1 log_10_ TCID_50_/d; 3–7 d) then rapidly (4 – 5 log_10_ TCID_50_/d; 7–9 d) [15]. We previously quantified this biphasic viral decline with a mathematical model, which indicated that the rate of infected cell clearance increases as the density of infected cells decreases [15]. The timing of the second, rapid viral decay phase coincides with the expansion of CD8^+^ T cells, which are the primary cell responsible for clearing infected cells and resolving the infection [20–24], and, to a lesser extent, neutralizing antibodies [21,22,24–26] and cytotoxic CD4^+^ T cells [27]. For the CD8^+^ T cell response, in particular, it remains unclear whether the efficacy of these cells is dictated by their own density [28,29], infected cell density [30–32], or both [33]. While quantifying dynamically changing CD8^+^ T cell efficacy is difficult *in vitro* and *in vivo*, the question is ripe for *in silico* investigation. Several modeling studies have described CD8^+^ T cell-mediated infected cell clearance for various viral infections, including IAV, HIV, and LCMV (e.g., as in [17,28,30,33–42]). Some of these studies have also attempted to link this response and others to inflammation or disease severity [39,40,43], but have had limited success even with substantial immunologic data. In addition, for IAV infections in particular, varied efficiency of the CD8^+^ T cell response throughout the course of infection, their early antigen-specific and lung-resident responses [44–46], and the balanced consequences on viral loads and host pathology have not yet been investigated in detail.

A better understanding of infected cell clearance may also yield insight into the damage induced to the lung and the ensuing immunopathology during IAV infection. In general, widespread alveolar disease is observed in patients who succumb to the infection [47], and CT scans show bilateral and multi-focal areas of damage (e.g. as in [48–50]). Further, hospitalized patients that died as a result from infection with the 2009 H1N1 influenza virus had large numbers of CD8^+^ T cells present in their lung tissue [7]. Large pulmonary populations of CD8^+^ T cells contribute to lung injury by targeting IAV-infected cells (reviewed in [5]) and inducing ‘bystander damage’ to uninfected epithelial cells [51]. However, their population levels do not necessarily indicate active infected cell removal or immunopathology as their extended presence in the lung also contributes to surveillance and repair [52,53]. Macrophages and neutrophils also persist in the lung following viral clearance, and their role in inflammation and tissue damage is well documented (reviewed in [9, 54]). A favorable outcome requires a balance between immune-mediated protection and pathology, and the viral-immunological landscape that drives disease and the marker that distinguish mild from severe influenza are complex. This is particularly true in humans [55] where obtaining high quality data from the lung is challenging, and viral loads in the upper respiratory tract do not reflect the lower respiratory tract environment.

The accumulation of damage to the epithelium during IAV infection, either from virally-induced cell lysis or immune-mediated effects, is relatively understudied. We and others have modeled the lung damage and inflammation inflicted during pulmonary infections (e.g., as in [40,56–59]) but did so without sufficient data to constrain the model formulations. The difficulty in measuring the dynamics of infected cells and in establishing how damage correlates to specific host responses has been the primary impediment. However, recent technological advances, including the use of reporter viruses [60–63] and lung histomorphometry [12, 64–67], have provided an opportunity to acquire these types of measurements. Whole lung histomorphometry, which is broadly defined as a quantitative microscopic measurement of tissue shape, has recently increased in use due to the ability to directly stain, visualize, and quantify areas of infected cells. Even with these techniques, quantitative data over the course of infection is not currently available. Having data such as these should help reconcile potential nonlinearities in infected cell clearance and provide insight into the accumulated lung inflammation and damage, which are generally thought to be markers of disease severity. In addition, it should bolster the development of robust, predictive computational models, which have historically lacked constraint to these types of data.

In general, disease severity does not seem to be directly correlated to viral loads or specific immunological components. In humans infected with IAV, viral loads typically increase prior to the onset of systemic symptoms, which peak around 2–3 d post-infection (pi) [18, 68, 69]. Some symptoms (e.g., fever) are cytokine mediated [70], but respiratory symptoms often last longer and can remain following viral clearance [68, 71]. This is particularly true when there is scarring of the lung tissue. In the murine model, weight loss is used as an indicator of disease progression and severity, where greater weight loss corresponds to more severe disease [72–74]. Animal weights generally drop slowly in the first several days during an IAV infection and more rapidly as the infection begins to resolve [19, 75]. This is unlike viral load dynamics in these animals, which increase rapidly during the first 0–3 d pi then remain relatively constant prior to resolution [15]. Because weight loss can occur following resolution of the infection, immune-mediated pathology is thought to play a role [5,74,76–78]. Host and pathogen factors, such as age, viral proteins, and inoculum size, can also influence disease progression [13, 14, 79, 80]. While the precise contribution of different factors to IAV-associated disease and mortality remain elusive, having tools that can link these with disease severity and decipher correlation from causation would improve our ability to effectively predict, understand, and treat the disease.

To gain deeper insight into the dynamics of viral resolution and investigate the connection between viral loads, damage, inflammation, and disease severity, we first simultaneously measured viral loads and CD8^+^ T cells daily from the lungs of BALB/cJ mice infected with influenza A/Puerto Rico/8/34 (H1N1) (PR8). We then developed a model that describes their kinetics to explore the mechanisms and dynamics of CD8^+^ T cell influx and their efficiency in removing virus-infected cells. The model agreed with our previous results that infected cells are cleared in a density-dependent manner, and showed accuracy in predicting the dynamics of effector and memory CD8 phenotypes. Our model predicted that infection duration is dependent on the magnitude of CD8^+^ T cells rather than their efficacy, which we validated by depleting CD8^+^ T cells. Further exploring these findings through quantitative whole lung histomorphometry corroborated the model-predicted infected cell dynamics and the direct, nonlinear relation between infected cell clearance and CD8^+^ T cell expansion. Using the inflammation scores, we developed an equation that relates these to the infected cell dynamics, and showed their distinction from lung injury and correlation to macrophages and neutrophils. In addition, these data revealed nonlinear connections between disease severity, the amount of the lung injured during infection, and inflammation. Our data, model, and analyses provide a robust quantification of the density-dependent nature of CD8^+^ T cell-mediated clearance, and the critical connections between these cells and the dynamics of viral loads, infected cells, lung injury, inflammation, and disease severity.

## 1 Results

### Virus and CD8^+^ T cell kinetics

In animals infected with 75 TCID_50_ PR8, virus rapidly infects cells or is cleared within 4 h pi and is undetectable in all animals at this time (Figure 1). Virus then increases exponentially and peaks after ~2 d pi. Following the peak, virus enters a biphasic decline. In the first phase (2–6 d pi), virus decays slowly and at a relatively constant rate (0.2 log_10_ TCID_50_/d) [15]. In the second phase (7–8 d pi), virus is cleared rapidly with a loss of 4 – 5 log_10_ TCID_50_ in 1–2 d (average of –3.8 log_10_ TCID_50_/d) [15]. CD8^+^ T cells remain at their baseline level from 0–2 d pi, where ~15% are located in the circulating blood, ~75% in the lung vasculature, and ~10% in the lung parenchyma (Supplementary file 1A) with the majority in the lung as CD43^−^CD127^+^. IAV-specific CD8^+^ T cells that are recently primed (CD25^+^CD43^+^) or effector cells (CD25^−^CD43^+^) [46] begin appearing in the lung and increase slightly from 2–3 d pi but remain at low levels. These cells are thought to proliferate within the lung at least once by 4 d pi [45], and their populations briefly contract (3–5 d pi) before expanding rapidly (5–8 d pi). During the expansion phase, >95% of recovered cells are in the lung (~70% in the parenchyma and ~30% in the vasculature) with CD25^+^CD43^+^ and CD25^−^CD43^+^ as the predominant phenotypes. This is in accordance with prior studies that showed these phenotypes are IAV-specific and that their expansion dynamics in the lung were not altered by removing blood-borne CD8^+^ T cells from the analysis [46,81].

**Figure 1.**
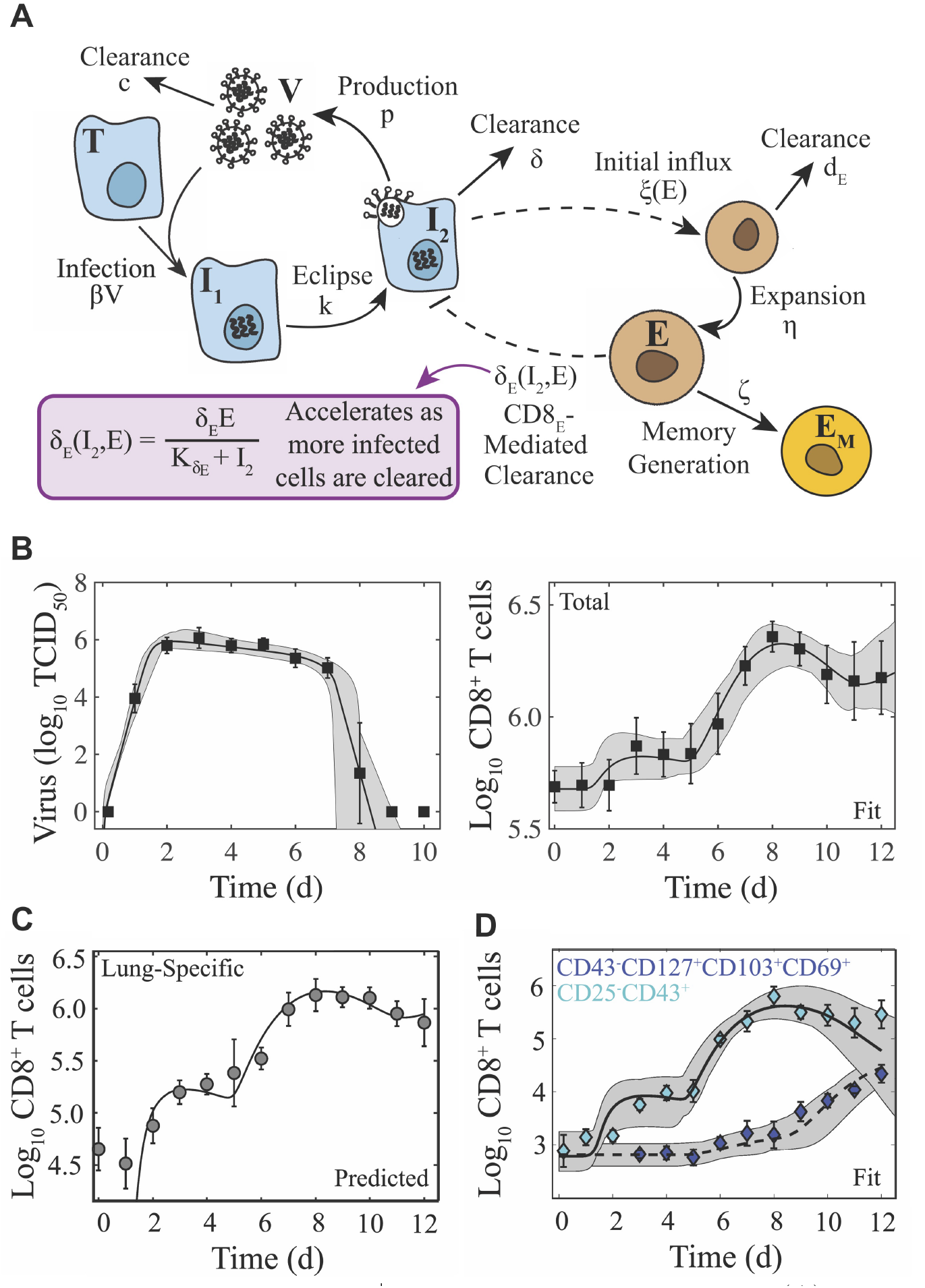
Schematic and fit of the CD8^+^ T cell viral kinetic model. (A) Schematic of the CD8^+^ T cell model in Equations (1)–(6). In the model, target cells (*T*) are infected at rate *βV*. Infected cells (*I*_1_) undergo an eclipse phase and transition to become productively-infected cells (*I*_2_) at rate *k*. Virus (*V*) is produced by infected cells at rate *p* and is cleared at rate *c*. Infected cells are cleared at rate *δ* by non-specific mechanisms and at rate *δ_E_*(*I*_2_,*E*) by effector CD8^+^ T cells (*E*; denoted CD8_E_). The dashed lines represent interactions between infected cells and CD8_E_. Initial CD8_E_ influx (*ξ*(*E*) = *ξ*/(*K_E_* + *E*)) is proportional to infected cells and is limited by CD8_E_ with half-saturation constant *K_E_*. CD8_E_ expansion (*η*) occurs proportional to infected cells *τ_E_* days ago. Memory CD8^+^ T cell (*E_M_*; denoted CD8_M_) generation occurs at rate *ζ* and proportional to CD8_E_ *τ_M_* days ago. (B) Fit of the CD8^+^ T cell model (Equations (1)–(6)) to virus and total CD8^+^ T cells from the lungs of mice infected with 75 TCID_50_ PR8 (10 mice per time point). The total number of CD8^+^ T cells is *Ê* = *E* + *E_M_* + *Ê*_0_. (C) Total CD8^+^ T cells in the lung parenchyma (gray circles) and overlay of the model predicted values (*E* + *E_M_*). (D) Fit of the model to virus, CD25^−^CD43^+^ CD8^+^ T cells (cyan diamonds; *E*), and CD43^−^CD127^+^ CD103^+^CD69^+^ CD8^+^ T cells (blue diamonds; *E_M_*) (5 mice per time point). The solid and dashed black lines are the optimal solutions and the gray shading is are the model solutions using parameter sets within the 95% CIs. Parameters are given in Table 1. Data are shown as mean ± standard deviation.

CD8^+^ T cell expansion corresponds to the second viral decay phase with sixty percent of mice clearing the infection by 8 d pi and the other forty percent by 9 d pi (Figure 1). Most CD43^+^CD8^+^ T cell phenotypes decline following viral clearance (8–10 d pi) but do not return to their baseline level by 12 d pi. Long-lived antigen-specific memory phenotypes down regulate CD43 [82–84] and gradually increase substantially beginning at 9 d pi with most as CD25^−^, which is qualitatively similar to other studies [83]. At 12 d pi, ~55% of CD8^+^ T cells remain in the lung parenchyma and ~20% in the circulating blood indicating exit from the lung (Supplementary file 1A).

### Viral kinetic model with density-dependent CD8^+^ T cell-mediated clearance

We previously described the viral load kinetics and biphasic decline using a density-dependent (DD) model in Equations (A9)–(A12) [15], which assumes that the rate of infected cell clearance increases as the density of infected cells decreases (i.e., *δ_d_*(*I*_2_) = *δ_d_*/(*K_δ_* + *I*_2_)). Because the rapid decay of virus is thought to be due to the clearance of infected cells by CD8^+^ T cells, it is unknown if early CD8^+^ T cell presence contributes to infected cell clearance, and no model has captured the entire CD8^+^ T cell time course, we developed a model that describes the dynamics of these cells and their efficiency in resolving the infection (Equations (1)–(6); Figure 1A). The model includes equations for effector (*E*, denoted CD8_E_) and memory (*E_M_*, denoted CD8_M_) CD8^+^ T cells, and two mechanisms of infected cell clearance. The first mechanism is from unspecified, innate mechanisms, which is relatively constant (*δ*) and primarily acts during the first viral decay phase (2–6 d pi). The second is the CD8_E_-mediated infected cell clearance, which occurs at a rate that increases as the density of infected cells decreases (*δ_E_*(*I*_2_, *E*) = *δ_E_E*/(*K_δ_E__* +*I*_2_)) and primarily acts during the rapid, second viral decay phase (7–8 d pi). Excluding this density dependence entirely resulted in a significant and premature decline in viral loads, which disagreed with the experimental data. We also tested whether the density dependence could be included in the CD8^+^ T cell expansion rate rather than in the infected cell clearance rate (see Equations (A1)–(A6)) as other models have done (e.g., as in [40–42]). This modification yielded a close fit to the CD8^+^ T cell data at 6–10 d pi but not at the early time points (Figure A1). In addition, the viral load data was underestimated at 7 d pi causing the solution to miss the rapid decline between 7–8 d pi and result in a statistically poorer fit. Thus, retaining the density-dependence in the rate of infected cell clearance most accurately captured the entire dataset. The model includes terms for the initial CD8_E_ influx at 2–3 d pi (*ξI*_2_/(*K_E_* + *E*)) and for CD8_E_ expansion (*ηEI*_2_(*t* – *τ_E_*)), which accounts for the larger increase between 5–8 d pi. To capture the contraction of CD8^+^ T cells between these times (3–5 d pi), it was necessary to assume that the initial CD8_E_ influx is regulated by their own population (i.e., *ξ*(*E*) = *ξ*/(*K_E_* + *E*)). In both terms, the increase is proportional to the number of infected cells. Although memory CD8^+^ T cells were not the primary focus here, it was necessary to include the CD8_M_ population because CD8^+^ T cells are at a significantly higher level at 10–12 d pi than at 0 d pi (Figure 1B). Fitting the model simultaneously to viral loads and total CD8^+^ T cells from the (non-perfused) lungs of infected animals illustrated the accuracy of the model (Figure 1B). The resulting parameter values, 95% confidence intervals (CIs), ensembles, and histograms are given in Table 1, in Figure 2, and in Supplementary file 1B.

**Figure 2.**
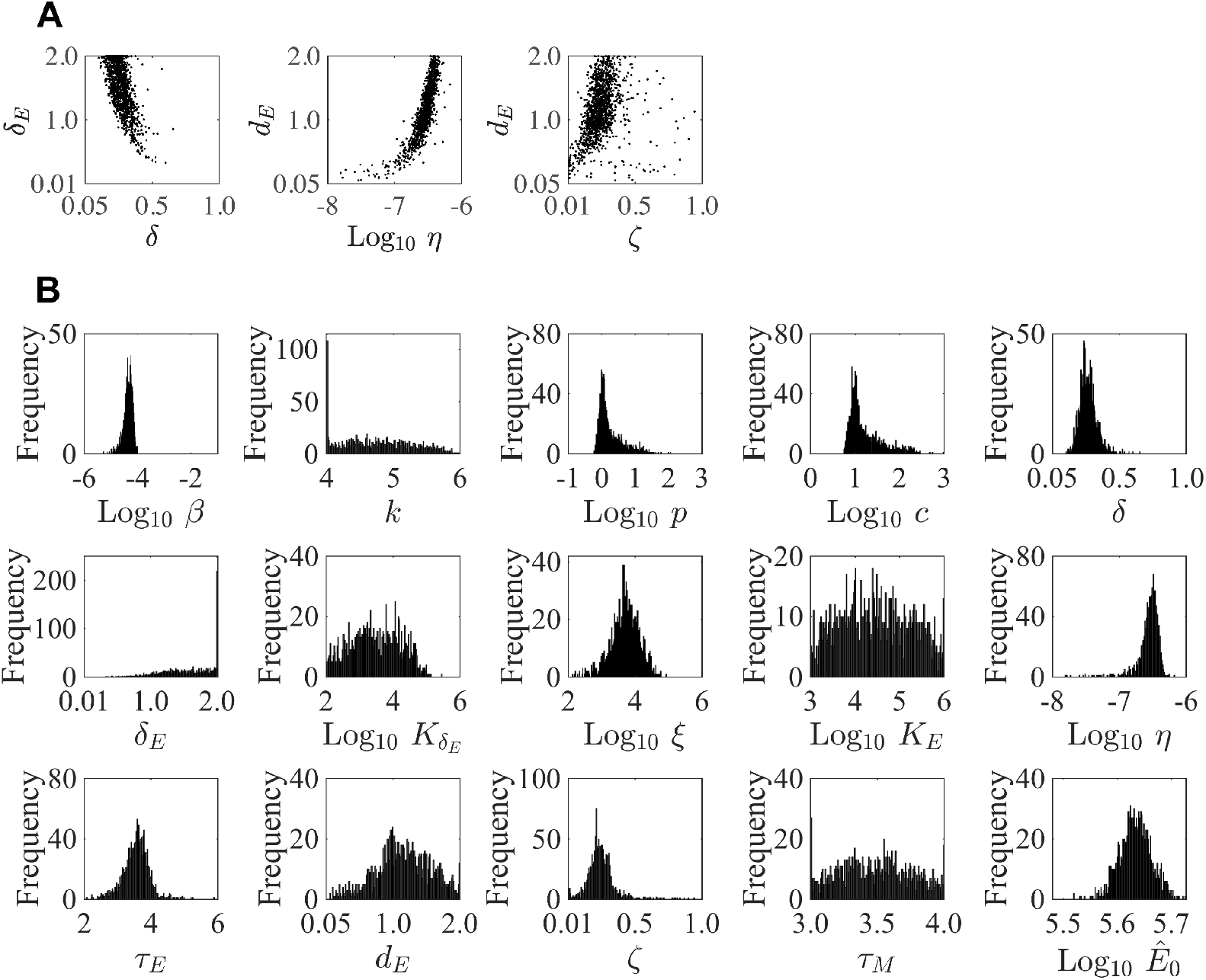
Parameter ensembles and histograms. Parameter ensembles (Panels A) and histograms (Panel B) resulting from fitting the CD8^+^ T cell viral kinetic model (Equations (1)–(6)) to viral titers and total CD8^+^ T cells from mice infected with 75 TCID_50_ PR8. (A) The rates of infected cell clearance by non-specific mechanisms (*δ*) and by CD8_E_ (*δ_E_*) are slightly negatively correlated. Correlations were also present between the rates of CD8_E_ clearance (*d_E_*), CD8_E_ expansion (*η*), and CD8_M_ generation (*ζ*). The axes limits reflect imposed bounds. Additional ensemble plots are in Supplementary file 1B. (B) The histograms show that the majority of parameters are well-defined with the exception of the eclipse phase transition rate (*k*), one of the half-saturation constants (*K_E_*), and the CD8_M_ generation delay (*τ_M_*).

**Table 1.** CD8^+^ T cell model parameters. Parameters and 95% confidence intervals obtained from fitting the CD8^+^ T cell model (Equations (1)–(6)) to viral titers and total CD8^+^ T cells (‘Total CD8’) or viral titers, CD25^−^CD43^+^CD8^+^ T cells, and CD43^−^CD127^+^CD103^+^CD69^+^CD8^+^ T cells (‘Specific CD8_E,M_ Phenotypes’) from mice infected with 75 TCID_50_ PR8. CD8_E_ and CD8_M_ denote effector (*E*) and memory (*E_M_*) CD8^+^ T cells, respectively. The total number of CD8^+^ T cells is *Ê* = *E* + *E_M_* + *Ê*_0_ and is denoted by CD8.

| Parameter | Description | Units | Total CD8 |  | Specific CD8<sub>E,M</sub> Phenotypes |  |
| --- | --- | --- | --- | --- | --- | --- |
|  |  |  | Value | 95% CI | Value | 95% CI |
| $\beta$ | Virus infectivity | TCID<sub>50</sub><sup>-1</sup> d<sup>-1</sup> | $6.2 \times 10^{-5}$ | $[5.3 \times 10^{-6}, 1.0 \times 10^{-4}]$ | $3.7 \times 10^{-5}$ | $[1.1 \times 10^{-5}, 9.4 \times 10^{-5}]$ |
| $k$ | Eclipse phase transition | d<sup>-1</sup> | 4.0 | [4.0, 6.0] | 5.1 | [4.0, 6.0] |
| $p$ | Virus production | TCID<sub>50</sub> cell<sup>-1</sup> d<sup>-1</sup> | 1.0 | $[5.8 \times 10^{-1}, 1.1 \times 10^2]$ | 1.5 | [0.73, 13.6] |
| $c$ | Virus clearance | d<sup>-1</sup> | 9.4 | $[5.6, 9.5 \times 10^2]$ | 12.1 | [5.8, 17.5] |
| $\delta$ | Infected cell clearance | d<sup>-1</sup> | $2.4 \times 10^{-1}$ | $[1.0 \times 10^{-1}, 6.6 \times 10^{-1}]$ | $3.0 \times 10^{-1}$ | $[1.9 \times 10^{-1}, 5.9 \times 10^{-1}]$ |
| $\delta_E$ | Infected cell clearance by CD8<sub>E</sub> | cells CD8<sub>E</sub><sup>-1</sup> d<sup>-1</sup> | 1.9 | $[3.3 \times 10^{-1}, 2.0]$ | 5.7 | [1.7, 8.5] |
| $K_{\delta_E}$ | Half-saturation constant | cells | $4.3 \times 10^2$ | $[1.0 \times 10^2, 2.9 \times 10^5]$ | $1.3 \times 10^2$ | $[1.0 \times 10^1, 8.6 \times 10^2]$ |
| $\xi$ | CD8<sub>E</sub> infiltration | CD8<sub>E</sub><sup>2</sup> cell<sup>-1</sup> d<sup>-1</sup> | $2.6 \times 10^4$ | $[1.3 \times 10^2, 8.7 \times 10^4]$ | $2.9 \times 10^3$ | $[8.0 \times 10^2, 1.7 \times 10^4]$ |
| $K_E$ | Half-saturation constant | CD8<sub>E</sub> | $8.1 \times 10^5$ | $[1.0 \times 10^3, 1.0 \times 10^6]$ | $2.2 \times 10^6$ | $[1.1 \times 10^6, 8.3 \times 10^6]$ |
| $\eta$ | CD8<sub>E</sub> expansion | cell<sup>-1</sup> d<sup>-1</sup> | $2.5 \times 10^{-7}$ | $[1.6 \times 10^{-8}, 6.7 \times 10^{-7}]$ | $3.5 \times 10^{-7}$ | $[2.3 \times 10^{-7}, 5.2 \times 10^{-7}]$ |
| $\tau_E$ | Delay in CD8<sub>E</sub> expansion | d | 3.6 | [2.1, 5.9] | 3.3 | [2.6, 3.8] |
| $d_E$ | CD8<sub>E</sub> clearance | d<sup>-1</sup> | 1.0 | $[5.1 \times 10^{-2}, 2.0]$ | 1.1 | $[3.3 \times 10^{-1}, 2.5]$ |
| $\zeta$ | CD8<sub>M</sub> generation | CD8<sub>M</sub> CD8<sub>E</sub><sup>-1</sup> d<sup>-1</sup> | $2.2 \times 10^{-1}$ | $[1.0 \times 10^{-2}, 9.4 \times 10^{-1}]$ | $3.2 \times 10^{-2}$ | $[1.0 \times 10^{-2}, 2.2 \times 10^{-1}]$ |
| $\tau_M$ | Delay in CD8<sub>M</sub> generation | d | 3.5 | [3.0, 4.0] | 3.3 | [2.1, 5.3] |
| $\hat{E}_0$ | Baseline CD8 or CD8<sub>E</sub> | CD8 or CD8<sub>E</sub> | $4.2 \times 10^5$ | $[3.3 \times 10^5, 5.3 \times 10^5]$ | $6.0 \times 10^2$ | $[3.1 \times 10^2, 1.8 \times 10^2]$ |
| $\hat{E}_{M0}$ | Baseline CD8<sub>M</sub> | CD8<sub>M</sub> | - | - | $6.5 \times 10^2$ | $[3.2 \times 10^2, 1.5 \times 10^3]$ |
| $T(0)$ | Initial uninfected cells | cells | $1 \times 10^7$ | - | $1 \times 10^7$ | - |
| $I_1(0)$ | Initial infected cells | cells | 75 | - | 75 | - |
| $I_2(0)$ | Initial infected cells | cells | 0 | - | 0 | - |
| $V(0)$ | Initial virus | TCID<sub>50</sub> | 0 | - | 0 | - |
| $E(0)$ | Initial CD8<sub>E</sub> | CD8<sub>E</sub> | 0 | - | 0 | - |
| $E_M(0)$ | Initial CD8<sub>M</sub> | CD8<sub>M</sub> | 0 | - | 0 | - |

Plotting the model predicted dynamics for the lung-specific CD8^+^ T cells (CD8_L_ = E + E_M_) illustrated the accuracy of the model in predicting their dynamics within the lung parenchyma without fitting to these data (Figure 1C). One benefit of using the total CD8^+^ T cells is that the model automatically deduces the dynamics of effector-mediated killing and memory generation without needing to specify which phenotypes might be involved as these may be dynamically changing. However, the rates of expansion, contraction, and infected cell clearance may be different if only certain phenotypes are engaged. Thus, we examined whether the model could fit the dynamics of the predominant effector (CD25^−^CD43^+^) and memory (CD43^−^CD127^+^CD103^+^CD69^+^) phenotypes. Re-fitting the model to these data suggested that no changes to model formulation were needed and there were only small alterations to select CD8-specific parameter values (Figure 1D; Table 1).

Plotting the model ensembles revealed a correlation between the two infected cell clearance parameters (*δ* and *δ_E_*; Figure 2B), which represent the efficacy of the non-specific immune response and the CD8^+^ T cell response, respectively. Performing a sensitivity analysis showed that the viral load dynamics do not change substantially when these parameters are increased or decreased (Figure A2). However, decreasing the rate of non-specific infected cell clearance (i.e., lower *δ*) resulted in a significant increase in the number of CD8_E_ due to the small increase in the number of infected cells (Figures A2–A3). Even with a larger CD8_E_ population, recovery was delayed by only ~ 0.1 d. Given the correlation between *δ* and *δ_E_* (Figure 2B), a more efficient CD8_E_ response (i.e., higher *δ_E_*) may be able to overcome this short delay in resolution. The lack of sensitivity to changes in the infected cell clearance parameters is in contrast to the DD model, where the viral dynamics were most sensitive to perturbations in *δ_d_* (Figure A2) [15], which encompasses multiple processes. With CD8^+^ T cells explicitly included in the model, the infection duration was most sensitive to changes in the rate of CD8_E_ expansion (*η*) (Figures A2, A3; discussed in more detail below).

Examining the parameter ensembles and sensitivity analysis also yielded insight into how other model parameters affect the CD8^+^ T cell response. The rates of CD8_E_ expansion (*η*) and clearance (*d_E_*) were slightly correlated, indicating a balance between these two processes (Figure 2A). This correlation and the sensitivity of *η* produced model dynamics that were also sensitive to changes in the CD8_E_ clearance rate (*d_E_*) (Figure A4). As expected, the rates of CD8_M_ generation (*ζ*) and CD8_E_ clearance (*d_E_*) were correlated (Figure 2A). It has been estimated that approximately 5–10% of effector CD8^+^ T cells survive to become a long-lasting memory population [85]. Despite the inability to distinguish between CD8_E_ and CD8_M_ in the total CD8^+^ T cell data, the model predicts that 17% of CD8_E_ transitioned to a memory class by 15 d pi. When considering only the CD25^−^CD43^+/-^ effector and memory phenotypes, the model estimates this value to be ~ 7%. Additional insight into the regulation of the CD8^+^ T cell response, results from the model fitting, and a comparison of the DD model and the CD8^+^ T cell model are included in Appendix A.2.

### Density-dependent infected cell clearance

Given the accuracy of the model, we next sought to gain further insight into the nonlinear dynamics of CD8^+^ T cell-mediated infected cell clearance. Plotting the clearance rate (*δ_E_*(*I*_2_,*E*) = *δ_E_E*/(*K_δ_E__* +1_2_)) as a function of infected cells (*I*_2_) and CD8_E_ (*E*) (Figure 3) confirmed that there is minimal contribution from CD8_E_-mediated clearance to viral load kinetics or infected cell kinetics prior to 7 d pi (Figure 3A–C, markers a-b). At the initiation of the second decay phase (7 d pi), the clearance rate is *δ_E_*(*I*_2_,*E*) = 3.5/d (Figure 3A–B, marker c). As the infected cell density declines towards the half-saturation constant (*K_δ_E__* = 4.3 × 10^2^ cells), the clearance rate increases rapidly to a maximum of 4830/d (Figure 3A–C, markers d–g). The model predicts that there are 6 × 10^5^ infected cells remaining at 7 d pi, which can be eliminated by CD8_E_ in 6.7 h.

**Figure 3.**
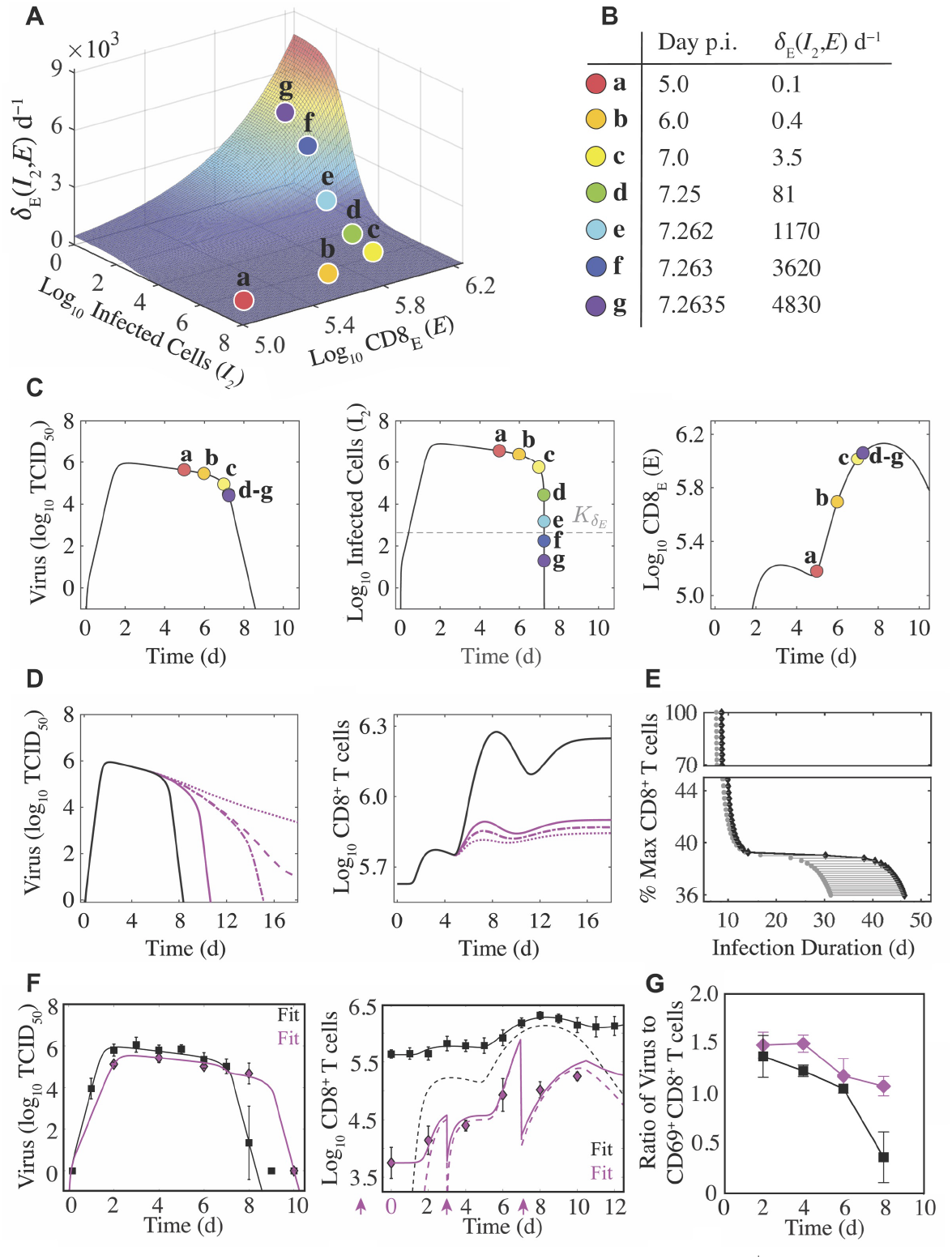
Density-dependent infected cell clearance by CD8^+^ T cells and their impact on recovery time. (A) The rate of CD8_E_-mediated infected cell clearance (*δ_E_* (*I*_2_, *E*) = *δ_E_E*/(*K_δ_E__* + *I*_2_)) plotted as a function of infected cells (*I*_2_) and effector CD8^+^ T cells (E; CD8_E_). The colored markers (denoted a–g) indicate the infected cell clearance rate that corresponds to different time points during the infection for the best-fit solution. (B) Values of *δ_E_* (*I*_2_, *E*) for the indicated time points associated with the markers a–g. (C) Corresponding locations of the various *δ_E_* (*I*_2_, *E*) values (markers a–g) on the best-fit solution of the CD8^+^ T cell model for virus (*V*), infected cells (*I*_2_), and CD8_E_ (*Ê*). (D) Solutions of the CD8^+^ T cell model (Equations (1)–(6)) for virus (*V*) and total CD8^+^ T cells (*Ê*) using the best-fit parameters (black line) and when varying the CD8_E_ expansion rate (*η*; magenta lines) to illustrate how different total CD8^+^ T cell magnitudes alter infection duration. The magenta lines are solutions from when the percent *Ê*_max_ relative to *Ê*_max_ from the best-fit solution was 42% (solid line), 39.2% (dash-dotted line), 39.1% (dashed line), or 37% (dotted line). (E) The time at which infected cells reach the half-saturation constant (*I*_2_ = *K_δ_E__*; gray circles) and the infection duration (time where log_10_ *V* = 0; black diamonds) are shown for the various CD8^+^ T cell magnitudes. The gray line between these points is the time required to eliminate *K_δ_E__* infected cells and achieve complete resolution of the infection (log_10_ *V* = 0). (F) Fit of the CD8^+^ T cell model (Equations (1)–(6)) to viral loads and CD8^+^ T cells (magenta diamonds) following depletion at −2, 0, 3, and 7 d pi (magenta arrows). The best model (see Supplementary file 1C) resulted in fewer target cells (*T*_0_), a lower CD8_E_ influx (*ξ*), and a higher CD8_E_ expansion rate (*η*). All other parameters were fixed to the best-fit value in Table 1. The solid lines are *Ê* = *E* + *E_M_* + *Ê*_0_ and the dashed lines are *E* for the cases where CD8s were depleted (magenta) and where they were not depleted (black). (G) Comparison of the log_10_ ratio of virus to CD69^+^CD8^+^ T cells with and without CD8^+^ T cell depletion (magenta and black, respectively). All data are shown as mean ± standard deviation.

To explore how recovery time is altered by varying CD8_E_ levels, we examined the resulting dynamics from increasing or decreasing the rate of CD8_E_ expansion (*η*). When *η* was increased by 50%, the CD8_E_ population increased by a substantial 670% (Figure A3). However, this was insufficient to significantly shorten the infection (8.4 d versus 7.8 d). The infection duration could be reduced if CD8_E_ expansion began earlier (i.e., smaller *τ_E_*; Figure A4). Although recovery is not significantly expedited by a larger CD8_E_ population, our model predicted that the infection would be dramatically prolonged if these cells are sufficiently depleted (Figures 3D–E and A3). This *in silico* analysis revealed a bifurcation in recovery time, such that the infection is either resolved within ~15 d pi or may last up to ~45 d if CD8_E_ are below a critical magnitude required to resolve the infection (Figure 3D–E). The critical number of total CD8^+^ T cells needed for successful viral clearance was 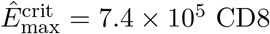, which was 39.2% of the maximum number of CD8^+^ T cells obtained from the best-fit solution (i.e., 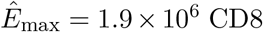). This corresponds to 17% of CD8_E_ (i.e., 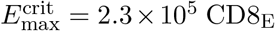; Figure 3D–E). The model analysis indicated that decreasing the total number of CD8^+^ T cells by as little as 0.1% from this critical level (i.e., 39.2% to 39.1%) lengthen the infection from 15 d pi to 25 d pi (Figure 3D).

### Dynamical changes during CD8^+^ T cell depletion

To further test our model formulation and identify how viral load kinetics are altered with dynamically changing CD8^+^ T cells, we infected groups of mice with 75 TCID_50_ and depleted these cells at −2 d, 0 d, 3 d, and 7 d pi with an anti-CD8*α* antibody (clone 2.43; Figure 3F). CD8^+^ T cells were reduced with over 99% efficiency and only 1.3% remained 2 d after depletion in the absence of infection (i.e., at 0 d pi; Figure 3F). CD8^+^ T cells remained > 1 log_10_ lower throughout the infection. Unexpectedly, the corresponding viral loads were significantly lower at 2 d pi (p=0.02) and consistently lower at other time points early in the infection (4 d pi (p=0.1) and 6 d pi (p=0.2)). As expected and predicted by our mathematical model, viral loads were significantly higher at 8 d pi (4.68 log_10_ TCID_50_ compared to 1.42 log_10_ TCID_50_; p=0.01). By 10 d pi, a sufficient number of CD8^+^ T cells were present and all animals had cleared the infection (Figure 3F). Interestingly, the number of CD8^+^ T cells at 10 d pi was only slightly higher than at 8 d pi (5.25 log_10_ versus 5.02 log_10_; p=0.064) further highlighting the density-dependent dynamics described above.

Given that viral loads were lower at early time points and that the anti-CD8*α* antibody is known to cause concentration-dependent changes in CD8_E_ differentiation, activation, and efficacy [86] in addition to resulting in death of the cells that would initiate activation of and removal by other immune cells, we did not expect our model to match the data without modulation of parameter values. However, we did expect that no changes to the model formulation would be required. In total, we tested >30 ‘models’ where 1-4 parameters were altered and found one model that was significantly better according to the AIC (Supplementary file 1C). In that model, the initial number of target cells (*T*_0_) was 2.5x lower (~ 4 × 10^6^ versus 1 × 10^7^ cells), the rate of initial CD8_E_ influx (*ξ*) was 2x lower (1.3 × 10^4^ versus 2.6 × 10^4^ CD8_E_^2^cell^−1^d^−1^), and the rate of CD8_E_ expansion (*η*) was 4x higher (1 × 10^−6^ versus 2.5 × 10^−7^ cell^−1^d^−1^). The second best model had a lower cost but was penalized by an additional parameter. That model suggested similar results but replaced the effect on *T*_0_ with a combination of a lower virus production rate (*p*) and higher non-specific infected cell clearance rate (*δ*). Both models resulted in fewer infected cells and, thus, have approximately equivalent interpretations.

To further examine these findings, we plotted the ratio of virus to activated (CD69^+^) CD8^+^ T cells (Figure 3G). At 2 d pi, the ratio was similar for the depleted and mock-treated groups (p=0.41) suggesting that the depletion-induced reduction in virus was proportional to the reduction in activated CD8^+^ T cells. However, at 4 d pi, the ratio in the depleted groups was significantly higher (p=0.0028) for otherwise similar levels of virus. This signifies that there were disproportionately low numbers of CD8^+^ T cells and, thus, reduced influx (i.e., lower *ξ*). By 6 d pi, the ratios were again similar (p=0.25) implying a higher expansion of these cells (i.e., higher *η*). Investigation into the predicted target cell reduction is included below.

### Modeling lung injury dynamics

To investigate the dynamics of infected cells and their clearance by CD8^+^ T cells, we quantified these cells and the progression and resolution of lung injury using serial whole lung histomorphometry (Figure 4A). Antigen-positive areas of the lung (“active” lesions) were first detectable at 2 d pi (Figure 4A–B), which coincides with the peak in viral loads. The infected area continued to increase in a nonlinear manner until 6 d pi, whereas viral loads remained high until 7 d pi (Figure 1B). At this time, resolution of the infection began and the infected area declined at a rate of ~28.7%/d between 6–7 d pi (Supplementary file 1D). Few to no infected cells were present at 8 d pi (Figure 4A). Correspondingly, virus was undetectable in most animals by 8 d pi (Figure 1B). Because the percent active lesion is a reflection of the influenza-positive cells, we examined whether the CD8^+^ T cell model accurately predicted these dynamics. In the model, the accumulated infection is defined by the cumulative area under the curve (CAUC) of the productively infected cells (*I*_2_). Plotting the percent active lesion against the CAUC of *I*_2_ shows that the model accurately reflects the cumulative infected cell dynamics and, thus, the infection progression within the lung (Figure 4B). Plotting the CAUC of *I*_2_ for all parameter sets in the 95% CIs and from fitting the model to specific phenotypes further illustrates the accuracy by showing that the heterogeneity in the histomorphometry data is captured (Figure 4B), which is larger than the heterogeneity in viral loads (Figure 1B). Plotting the predicted active lesion dynamics for the model parameterized to the data where CD8^+^ T cells were depleted suggested that there was a ~22% reduction in the active lesion Figure 4B.

**Figure 4.**
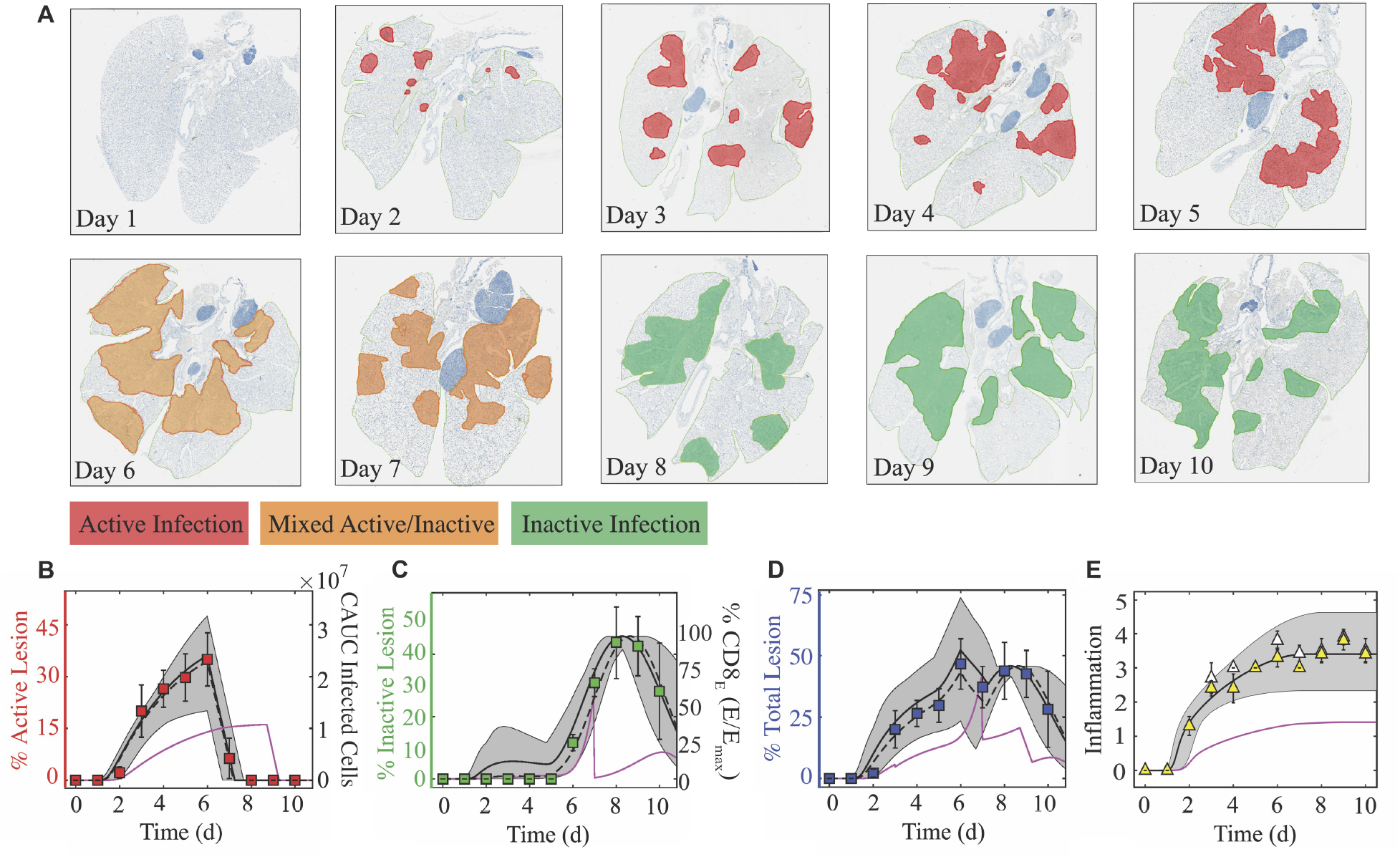
Histomorphometry and inflammation of the IAV-infected lung. (A) Whole lung sections with histomorphometry showing the areas of influenza NP-positive “active” lesions (red), previously infected “inactive” lesions with minimal antigen-positive debris (green), or mixed active and inactive regions (orange) throughout the infection. (B) Percent active lesion (red squares) plotted together with the cumulative area under the curve (CAUC) of the predicted infected cell dynamics (*I*_2_) obtained from fitting the CD8^+^ T cell model. The linear decline in the active lesion (−28.7%/d; see Supplementary file 1D) was used to estimate the decline after 6 d pi. (C) Percent inactive lesion (green squares) plotted together with the percent maximum CD8_E_ (*E/E*_max_) obtained from fitting the CD8^+^ T cell model. (D) The total lesion (blue squares) is the addition of the active and inactive lesions. To include all measurements on the same scale, the CAUC of *I*_2_ was multiplied by a scaling factor of 14.2% per 1 × 10^7^ cells, and the percent maximum CD8_E_ was multiplied by a scaling factor of 0.46%. (E) Fit of Equation (7) to the alveolar (white triangles) and interstitial (yellow triangles) inflammation scores. The solid black, dashed black, and solid magenta lines are the curves generated using the best-fit parameters obtained from fitting the model to the total CD8^+^ T cells, the CD25^−^CD43^+^ and CD43^−^CD127^+^ CD103^+^CD69^+^ CD8^+^ T cells, and the total CD8^+^ T cells during CD8 depletion, respectively. The gray shading are the curves generated using the 95% CI parameters from fitting the model to the total CD8^+^ T cells.

Antigen-negative, previously-infected or damaged areas of the lung (“inactive” lesions) are evident beginning at 5 d pi (Figure 4A,C). This resolution of the infection continued from 5–8 d pi, causing a 15.1%/d increase in the area of inactive lesions (Supplementary file 1D). Following this, healing of the injured lung is apparent as the inactive lesioned area declines (−14.5%/d from 9–10 d pi; Figures 4C and Supplementary file 1D). These dynamics generally parallel the CD8^+^ T cell dynamics but are nonlinearly correlated (Supplementary file 1D). Fitting a line to the CD8^+^ T cell data from 5–8 d pi indicated that the influx rate of all phenotypes is 4.94 × 10^5^ cells/d, of lung-specific phenotypes is 3.97 × 10^5^ cells/d, and of the CD25^−^CD43^+^ effector phenotype is 1.98 × 10^5^ cells/d (Supplementary file 1D). Thus, the model estimates that, on average, every 100,000 total, lung-specific, or CD25^−^CD43^+^ CD8^+^ T cells clear ~ 3.1%, ~ 3.8%, or ~ 7.6% of the lung, respectively. During the CD8^+^ T cell contraction phase, a similar linear regression analysis suggested that these cells decline at rates of ~ 4.13 × 10^5^ CD8/d (total), ~ 2.35 × 10^4^ CD8/d (lung-specific), and ~ 3.83 × 10^4^ CD8/d (CD25^−^CD43^+^) (Supplementary file 1D). Similar to the relation discussed above, the dynamics of the damaged areas of the lung corresponded to the dynamics of the percent maximum CD8_E_ (i.e., *E/E*_max_) in the model (Figure 4C). Our model suggested a ~37% reduction in the inactive lesion during CD8^+^ T cell depletion (Figure 4C). Adding the predicted dynamics for the active and inactive lesions agrees with the dynamics of the total lesion, but is slightly underestimated when using the model fit to CD25^−^CD43^+^ CD8^+^ T cells (Figure 4D). In addition, the predicted total lesion was reduced for the case where CD8^+^ T cells were depleted (Figure 4D).

### Modeling lung inflammation dynamics

In addition to measuring virus-induced lung damage, lung inflammation was scored (Figure 4E). Both alveolar and interstitial inflammation begin to increase at 2 d pi with the sharpest increase between 1–3 d pi. Inflammation continues to increase until 6 d pi with a maximum score of 3.5-4.0 out of 5. Resolution of inflammation was not evident during the time course of our data, which concluded at 10 d pi. This is in contrast to the lung damage inflicted by the virus, which begins declining at 8 d pi and shows that ~15% of the damage was repaired by 10 d pi. Inflammation was nonlinearly correlated to inflammatory macrophages and linearly correlated to neutrophils (Supplementary file 1E), which were excluded from the model. However, the knowledge that the model’s predicted infected cell dynamics are accurate and that these cells can produce cytokines and chemokines that attract cells like macrophages and neutrophils suggested that our model could be used to estimate inflammation. To do this, we fit Equation (7) to the inflammation scores while keeping all other parameters fixed to their best-fit values (Table 1). The results suggested that the equation captured the inflammation dynamics, and that the contribution from the initial infected cell class (*I*_1_) was *α*_1_ = 4.27 per 10^7^ cells/d and the contribution from the productively infected cells (*I*_2_) was *α*_2_ = 0.87 per 10^7^ cells/d. For the case where CD8^+^ T cells were depleted, Equation (7) estimated the inflammation scores to be reduced to ~1.5 (Figure 4E).

### Weight loss relates nonlinearly to lung injury and inflammation

To monitor disease progression, weight loss was measured daily throughout the course of infection (Figure 5). During the first 5 d pi, animals gradually lost ~4% of their initial weight. This was followed by a sharper drop (8%) at 6 d pi. Animal weights increased slightly at 7 d pi (~6%) before reaching peak weight loss (10–14%) at 8 d pi. Following virus resolution, the animals’ weights began to restore as the inactive lesions resolved (9–10 d pi; Figure 5A). Plotting weight loss against the percent total (active and inactive) lesioned area of the lung shows the similarity in their dynamics (Figure 5A) and revealed the nonlinear relation (Figure 5B). To further quantify their relationship, we fit the saturating function 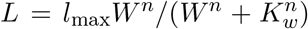 to these data, where *L* is the percent total lesioned area, *W* is percent weight loss, *l*_max_ is the maximum rate of the interaction, *K_w_* is the half-saturation constant, and *n* is the Hill coefficient. This function provided a close fit to the data (*R*^2^ = 0.92; Figure 5B) with best-fit parameters *l*_max_ = 39.7% total lesioned area, *K_w_* = 2.58% weight loss, and *n* = 5.24. Weight loss was also nonlinearly correlated to the alveolar and interstitial inflammation scores (Figure 5C), although the relation was slightly different compared to that for the lung injury data. Fitting the same function to these data independently for alveolar and interstitial inflammation also provided a close fit (*R*^2^ = 0.98 and *R*^2^ = 0.97; Figure 5D). Best-fit parameters for alveolar inflammation were *l*_*a*_max__ = 3.63 score, *K_a_w__* = 1.95% weight loss, and *n_a_* = 3.65. Best-fit parameters for interstitial inflammation were *l*_*i*_max__ = 3.40 score, *K_i_w__* = 1.96% weight loss, and *n_i_* = 3.15.

**Figure 5.**
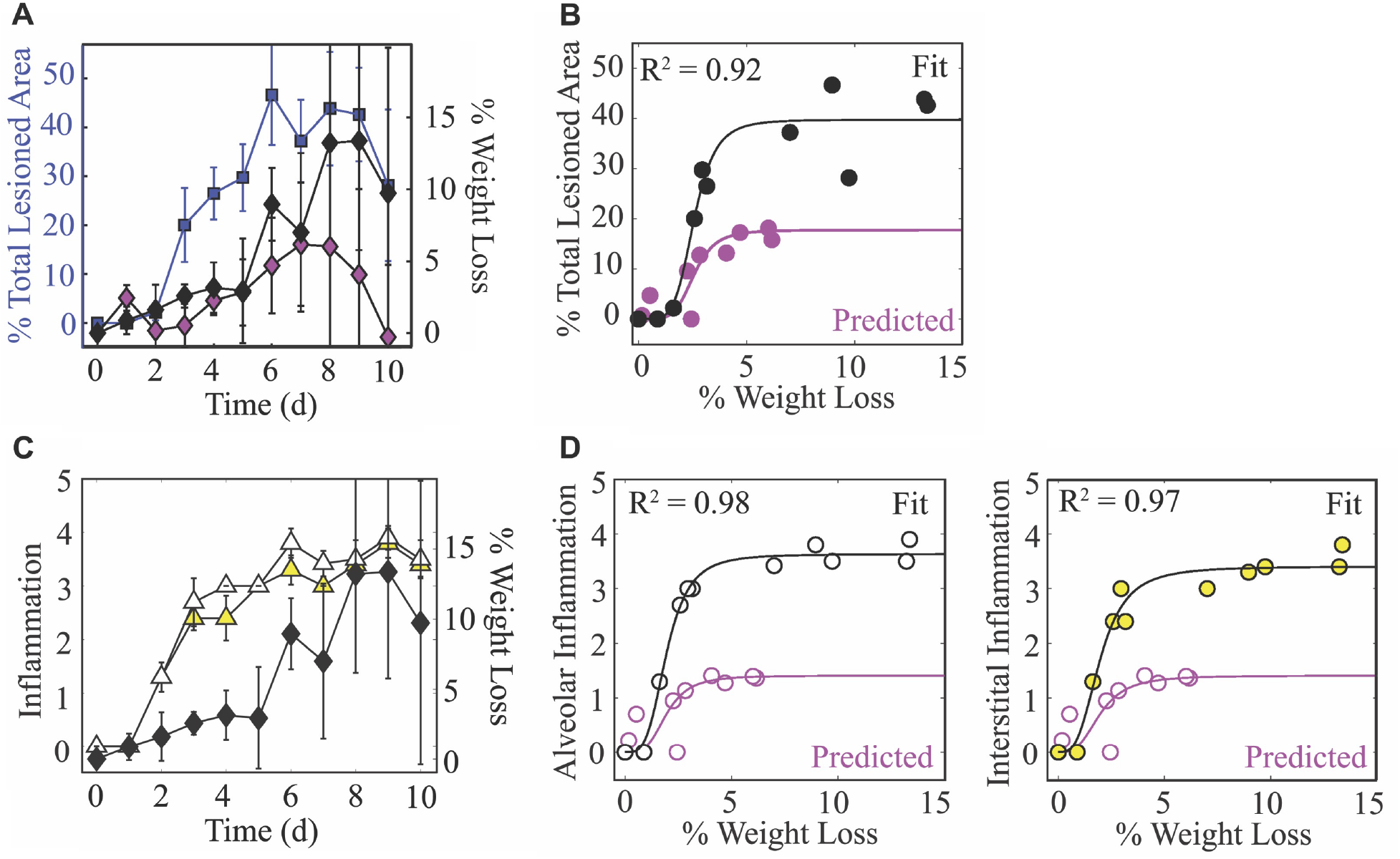
Weight loss dynamics and its relation to lung injury and inflammation. (A) The percent total (active and inactive) lesion (blue squares) plotted together with the percent weight loss (black diamonds) to illustrate their similar dynamics. (B) Fit of a saturating function 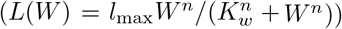 to the mean percent total lesioned area (L) and mean weight loss (*W*) for all time points. The best-fit parameters were *l*_max_ = 39.7% total lesioned area, *K_w_* = 2.6% weight loss, and *n* = 5.2. (C) The alveolar (white triangles) and interstitial inflammation (yellow triangles) plotted together with the percent weight loss (black diamonds). (D) Fit of a saturating function to the mean inflammation scores and mean weight loss for all time points. The best-fit parameters for alveolar inflammation were *l*_*a*_max__ =3.63 score, *K_aw_*=1.95% weight loss, and *n_a_* = 3.65, and for interstitial inflammation were *l*_*i*_max__ =3.40 score, *K_i_w__* =1.96% weight loss, and *n_i_* = 3.15.

## Discussion

Influenza A virus infections pose a significant threat to human health, and it is crucial to understand and have tools that can predict how the virus spreads within the lower respiratory tract, how specific immune responses contribute to infection control, and how these relate to disease progression. Although it has been difficult to directly relate these features and obtain high quality data from the lower respiratory tract in humans, we circumvented the challenge by pairing comprehensive experimental data with robust mathematical models and analyses. Our iterative model-driven experiment approach [87, 88] revealed important dynamic relations between virus, infected cells, immune cells, lung damage, inflammation, and disease severity (summarized in Figure 6). Identifying these nonlinear connections allows for more accurate interpretations of viral infection data and significant improvement in our ability to predict disease severity, the likelihood of complications, and therapeutic efficacy.

**Figure 6.**
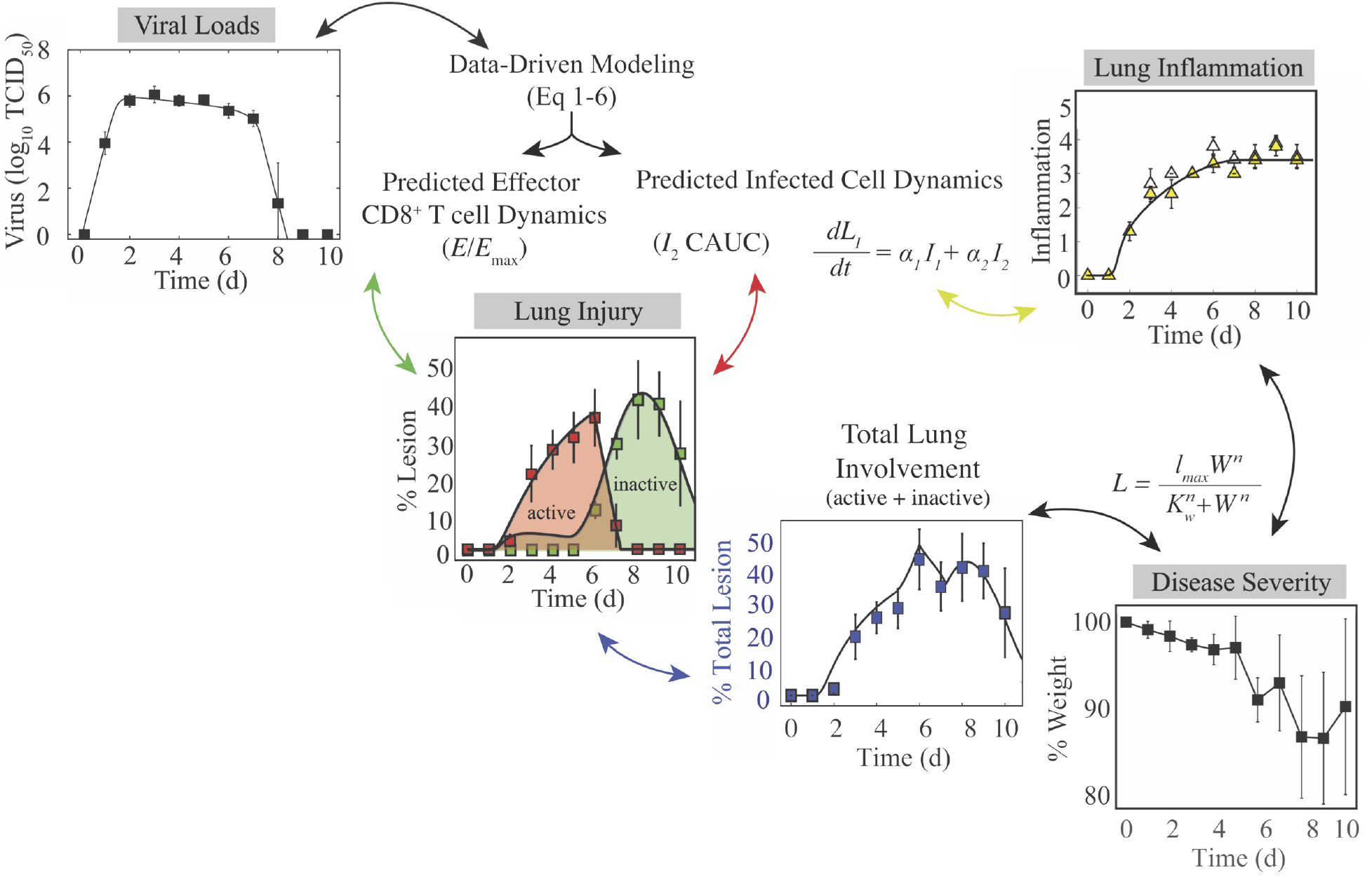
Summary of the connections between the kinetics of virus, infected cells, CD8^+^ T cells, lung injury, lung inflammation, and disease severity. Summary of the relations between the dynamics of virus, infected cells, CD8^+^ T cells, lung lesions, lung inflammation, and weight loss established by our analysis. Given that viral loads and weight loss are the most easily measured variables, our mathematical model (Equations (1)–(6)) could be used to estimate the kinetics of infected cells, CD8^+^ T cells, and inflammation. The cumulative area under the curve (CAUC) of the productively infected cell dynamics (I_2_) yields an estimate of the percent lung infected (active lesion) while the predicted relative CD8_E_ dynamics (*E/E_max_*) yields an estimate of the percent lung resolved (inactive lesion). The total amount of lung involved (% lung infected and % lung resolved) or the inflammation scores can then be used to estimate weight loss through the functions *L_l_*(*W*) and *L_I_*(*W*). These connections could be reversed and weight loss used to estimate viral load kinetics.

Our histomorphometric data and analyses provided a robust quantification of the extent of influenza virus infection within the lung and the dynamics of lung injury and inflammation. The images look strikingly similar to CT scans from infected patients, which also show extensive bilateral damage [48–50]. The model’s 95% CI predictions of the lesioned area nicely captured the heterogeneity in the data (Figure 4), which is remarkable given the minimal variability in the virus and CD8^+^ T cell data and model fit (Figure 1). The differential heterogeneity suggests that small changes to viral loads can induce significant changes in disease severity. Indeed, this has been observed when antivirals are administered [89–94] or when host responses are modulated [95–97]. It may also indicate that viral loads are not a reliable measure of infection severity. This knowledge should also aid future studies seeking to estimate therapeutic efficacy or the effects of immune modulation. There was a slight over-estimation of the inactive lesion early in the infection (1–5 d pi, dark gray shading; Figure 4) when using the fit to the total CD8^+^ T cells, which was abrogated when using the CD25^−^CD43^+^CD8^+^ T cells. However, these cells underestimated the total lesion at 6-7 d pi, suggesting contributions from other CD8^+^ phenotypes. In general, modeling different CD8^+^ T cell phenotypes may help clarify their *in vivo* functions and better define the rates of CD8-mediated infected cell clearance, but our results suggest that this is not necessarily required.

Our data and analyses provided strong support that the infected cell density impacts the rate at which they are cleared by effector CD8^+^ T cells (CD8_E_) (Figures 3–4, Supplementary file 1D). Although CD8^+^ T cell dynamics can be somewhat replicated when assuming the density dependence lies within their expansion, that assumption could not recover the viral load dynamics (Figure A1). Discriminating between these mechanisms is difficult *in vivo*, but the ability of the model in Equations (1)–(6) to capture the entire time course of CD8_E_ dynamics in multiple scenarios is compelling. Regardless of the mechanism, we first detected this density-dependence in a model that excluded specific immune responses (i.e., Equations (A9)–(A12)) [15]. Simple models like that one are useful to capture nonlinearities, but they cannot distinguish between different mechanisms.

Several factors may contribute to the density-dependent change in the rate of CD8_E_-mediated clearance. One possibility is that the slowed rate of clearance at high infected cell densities is due to a “handling time” effect, which represents the time required for an immune cell to remove an infected cell (e.g., as in [15,28,33,98–101]). When CD8_E_ interact with infected cells, a complex is formed for ~20–40 min [36,102–105]. Because CD8_E_ could not interact with other infected cells during this time, the global rate of infected cell clearance would be lowest when infected cells outnumber CD8_E_. In addition, contact by a single CD8_E_ may be insufficient to remove an infected cell [32]. Infected cell clearance is more frequently observed after interactions with multiple CD8_E_, with an average of 3.9 contact events either serially or simultaneously [32]. Thus, the high density of infected cells early in the infection reduces the probability that a single virus-infected cell would be targeted a sufficient number of times to induce cell death. However, as CD8_E_ accumulate and the density of infected cells decreases (Figure 3A), the probability of simultaneous interactions will increase. These interactions may also be influenced by CD8_E_ movement, where their velocity slows at the height of infected cell clearance (6–8 d pi) [52, 106]. This should reduce the time required to remove an infected cell and, thus, result in a higher efficiency. Moreover, it is possible that spatial limitations also contribute, such that a high infected cell density may hinder CD8^+^ T cells from reaching infected cells on the interior of the infected tissue. Crowding of immune cells at the periphery of infected lesions has been observed in other infections (e.g., in tuberculosis granulomas [107, 108]) and has been suggested in agent-based models of influenza virus infection [109].

The new knowledge about how the extent of infection within the lung also uncovered the nonlinear connection between disease severity and various measures of lung pathology (Figure 5). This discovery is significant because it suggests there is a unique link to both the extent of lung injury and inflammation, which are often used interchangeably yet have distinct dynamics (Figure 4) and correlate to different immune responses (Supplementary file 1F, Supplementary file 1G). In the same vein, both measurements can be estimated using the dynamics of infected cells (Figure 4) but with independent mathematical relations (CAUC of *I*_2_ versus Equation (7)). Our CD8 depletion data (Figure 3) and other experimental studies using histomorphometry [12] corroborate a relationship between weight loss and lung pathology during IAV infection. For example, animals treated with antivirals in various conditions (single or combination therapy and in immunocompetent or immunosuppressed hosts) demonstrated that, although viral loads are not always significantly reduced, the antiviral-induced reductions in weight loss were paired with decreased infected areas of the lung [12, 64]. Further examining lung injury and inflammation kinetics and their connection to weight loss in various experimental infection settings (e.g., different ages or sex, under therapy, other strains or viruses, etc.) and deciphering how innate responses contribute to these measurements should improve our predictive capabilities. This may be particularly helpful in understanding the exacerbated morbidity and mortality in elderly individuals, where there are lower viral loads, increased weight loss, symptoms, and/or mortality within animal models [13, 14].

This work demonstrates a significant potential for the easily obtained weight loss data to be used, analyzed, and interpreted within both modeling and experimental studies. In our data and others’ animal data, there is a spike in weight loss, symptom score, and/or inflammation at ~6 d pi [75, 79, 110, 111] (Figure 5). Our data suggests that this may be due to CD8_E_ activity while the infection continues to spread. The subsequent increase in animal weights starting at 7 d pi is concurrent with the decline in infected lesions (Figure 5). Interestingly, there was no corresponding decrease in inflammation, and the spike was not apparent in the CD8 depletion data where severity was reduced. The subsequent increase in weight loss increase following infection resolution (8–9 d pi; Figure 4C) could be attributed to CD8^+^ T cell-mediated pathology and/or to ongoing inflammation. The tighter correlation to inflammation might suggest the latter; however, some studies indicate that large numbers of CD8^+^ T cells pose a risk of acute lung tissue injury [5–9]. According to our model predictions, excessive CD8^+^ T cell numbers may augment disease progression yet do not improve recovery time (Figure A3). Instead, an earlier onset of CD8_E_ proliferation (i.e., smaller *τ_E_*) would be required to significantly shorten the infection (Figure A4). This aligns with evidence that hosts with adaptive immune responses primed by vaccine or prior infection recover more rapidly [112, 113]. While higher CD8^+^ T cell numbers have little impact on viral kinetics, our model and data agree with clinical and experimental studies from a wide range of host species that impaired CD8^+^ T cell responses can prolong an IAV infection [17,22,26,114–116]. In some scenarios, virus can persist for up to several weeks if CD8^+^ T cell-mediated clearance is unsuccessful (Figure 3; [26, 117, 118]). The bifurcation in recovery time revealed by the model suggests that this may occur when the number of lung-specific CD8^+^ T cells are less than 2.3 × 10^5^ cells (Figure 3D–E). Our CD8 depletion data show that the precise number will be dependent on other infection variabless. Minimally, we would expect this number to vary depending on parameters like the dose, rate of virus replication, and/or the infected cell number and lifespan, which has been noted in another modeling study that detailed similar bifurcating behavior [119]. Although some previously published models also suggested delayed resolution with depleted CD8^+^ T cell responses [17,37,38,120], this bifurcation has not been observed and their estimated delays in recovery do not amount to the long-lasting infections in immunodeficient patients [26, 117, 118]. Our model’s ability to capture the dynamics when CD8^+^ T cells are depleted is encouraging, and the data are an important reminder that experimental modulation of cell populations (e.g., through depletion or genetic knockouts) is complicated and that the data from such systems should be interpreted cautiously. It also brings into question studies that have used these types of data without validated mathematical models or quantification of other immunological variables.

In addition to illuminating the connections between viral kinetics and pathology, the histomorphometric data validated the model’s infected cell dynamics (Figure 4B). The dynamics of susceptible and infected cells throughout the infection and the accuracy of the target cell limited approximation used within influenza viral kinetic models have been questioned for several years [88, 121–126]. The ability of the model to accurately predict the histomorphometry and CD8^+^ T cell depletion data corroborates the use of this approximation, which assumes a limited number of available target cells and describes their decline by infection only. Although the slowing of viral loads beginning at 2 d pi could be due to a variety of innate immune-mediated mechanisms (e.g., macrophages, neutrophils, type I interferons, etc.), adding more complex dynamics to our model was unnecessary to describe the data. Further, the model and data agree that there are few infected cells during the time when viral loads are growing most rapidly (0–2 d pi; Figures 1B and 4B). We previously used this information to derive approximations for the model and gain deeper insight into how each parameter influences the kinetics [127], which has helped numerous studies interpret their results [15,17,128,129]. The data also supports the model’s hypothesis that that there is minimal clearance of infected cells prior to CD8_E_ expansion (Figure 4). The knowledge of the model’s accuracy and of the spatiality in the lung should aid investigation into the mechanisms that limit virus growth during the early stages of the infection.

Employing targeted model-driven experimental designs to examine and validate theoretical predictions like the ones presented here is pivotal to elucidating the mechanisms of infection spread and clearance [87, 88]. Examining other infections (e.g., coronaviruses), modifications to the dynamics (e.g., lethal doses), and the connection between lung measurements and those more easily acquired from upper respiratory tract will help refine the dynamical links between virus, host immune responses, and disease severity and identify their generalizability. Determining the factors that influence disease severity is vital to understanding the disproportionate mortality in at-risk populations (e.g., elderly) and to improving therapeutic design. This is particularly important because current antivirals alleviate symptoms but do not always effectively lower viral loads [89–94]. The predictive capabilities of validated models like the one here should prove useful in forecasting infection dynamics for a variety of scenarios. These tools and analyses provide a more meaningful interpretation of infection data, new ways to interpret weight loss data in animal models, and a deeper understanding of the progression and resolution of the disease, which will undoubtedly aid our ability to effectively combat influenza.

## 2 Methods and Materials

### Ethics statement

All experimental procedures were performed under protocols O2A-020 or 17-096 approved by the Animal Care and Use Committees at St. Jude Children’s Research Hospital (SJCRH) or the University of Tennessee Health Science Center (UTHSC), respectively, under relevant institutional and American Veterinary Medical Association (AVMA) guidelines. All experimental procedures were performed in a biosafety level 2 facility that is accredited by the American Association for Laboratory Animal Science (AALAS).

### Mice

Adult (6 week old) female BALB/cJ mice were obtained from Jackson Laboratories (Bar Harbor, ME) or Charles River Laboratories (Wilmington, Massachusetts). Mice were housed in groups of 5 mice in high-temperature 31.2cm × 23.5cm × 15.2cm polycarbonate cages with isolator lids (SJCRH) or in 38.2cm × 19.4cm × 13.0cm solid-bottom polysulfone individually ventilated cages (UTHSC). Rooms used for housing mice were maintained on a 12:12-hour light:dark cycle at 22 ± 2°C with 50% humidity in the biosafety level 2 facility at SJCRH (Memphis, TN) or UTHSC (Memphis, TN). Prior to inclusion in the experiments, mice were allowed at least 7 d to acclimate to the animal facility such that they were 7 weeks old at the time of infection. Laboratory Autoclavable Rodent Diet (PMI Nutrition International, St. Louis, MO; SJCRH) or Teklad LM-485 Mouse/Rat Sterilizable Diet (Envigo, Indianapolis, IN; UTHSC) and autoclaved water were available ad libitum. All experiments were performed under an approved protocol and in accordance with the guidelines set forth by the Animal Care and Use Committee at SJCRH or UTHSC.

### Infectious agents

All experiments were done using the mouse adapted influenza A/Puerto Rico/8/34 (H1N1) (PR8).

### Infection experiments

The viral infectious dose (TCID_50_) was determined by interpolation using the method of Reed and Muench [130] using serial dilutions of virus on Madin-Darby canine kidney (MDCK) cells. Mice were intranasally inoculated with 75 TCID_50_ PR8 diluted in 100μl of sterile PBS. In a subset of animals, CD8^+^ T cells were depleted by IP injection at days −2, 0, 3, and 7 pi with 100μg of the rat anti-CD8*α* antibody (clone 2.43) that was purified from ATCC hybridoma (per manufacturer instructions) in 250μl of PBS. Depletion efficiency was confirmed in the lung and spleen as described below. Mice were weighed at the onset of infection and each subsequent day to monitor illness and mortality. Mice were euthanized if they became moribund or lost 30% of their starting body weight. For viral load and total CD8^+^ T cell quantification, experiments were repeated three times and in each facility to ensure reproducibility and two complete experiments (ten animals per time point) were used for these studies. For all other experiments, the data was compared to prior results in addition to being repeated at select time points to ensure reproducibility, and one complete experiment (five animals per time point) was used. For pathology scoring and histomorphometry quantitation, five animals per time point were used. Power was calculated using G*Power.

### Lung harvesting for viral and cellular dynamics

For total CD8^+^ T cell quantification, mice were euthanized by CO_2_ asphyxiation. To distinguish blood-borne CD8^+^ T cells from those in the lung parenchyma, deeply anesthetized mice (5% isoflurane) were retro-orbitally injected with 3 μg of anti-CD45 antibody (PerCP, clone 30-F11, Biolegend) 3 min prior to euthanasia [46,81]. Mice were then euthanized by 33% isoflurane inhalation and their lungs perfused with 10 ml PBS prior to removal. For all experiments, lungs were then aseptically harvested, washed three times in PBS, and placed in 500μl sterile PBS. Whole lungs were digested with collagenase (1mg/ml, Sigma C0130), and physically homogenized by syringe plunger against a 40μm cell strainer. Cell suspensions were centrifuged at 4°C, 500xg for 7 min. The supernatants were used to determine the viral titers (TCID_50_) by serial dilutions on MDCK monolayers. Following red blood cell lysis, cells were washed in MACS buffer (PBS, 0.1M EDTA, 0.01M HEPES, 5mM EDTA and 5% heat-inactivated FBS). Cells were then counted with trypan blue exclusion using a Cell Countess System (Invitrogen, Grand Island, NY) and prepared for flow cytometric analysis as indicated below.

### Lung titers

For each animal, viral titers were obtained using serial dilutions on MDCK monolayers and normalized to the total volume of lung homogenate supernatant. The resulting viral loads are shown in Figure 1B and were previously published and utilized for calibration of the density-dependent model (Equations (A9)–(A12)) [15].

### Flow cytometric analysis

Flow cytometry (LSRII, BD Biosciences, San Jose, CA (SJCRH) or ZE5 Cell Analyzer, Bio-Rad, Hercules, CA (UTHSC)) was performed on the cell pellets after incubation with 200μl of a 1:2 dilution of Fc block (human-γ globulin) on ice for 30 min, followed by viability (Biolegend, Zombie Violet Fixable Viability Kit) and surface marker staining with anti-mouse antibodies. For total CD8^+^ T cell, macrophage, and neutrophil quantification, we used antibodies CD3e (BV786, clone 145-2C11, Biolegend), CD4 (PE-Cy5, clone RM4-5, BD Biosciences), CD8*α* (BV605, clone 53-6.7, BD Biosciences), Ly6C (APC, clone HK1.4, eBioscience), F4/80 (PE, clone BM8, eBioscience), CD11c (eFluor450, clone N418, eBioscience), CD11b (Alexa700, clone M1/70, BD Biosciences), MHC-II (FITC, clone M5/114.15.2, Invitrogen), CD49b (APCe780, clone DX5, eBioscience), Ly6G (PerCp-Cy5.5, clone 1A8, Biolegend). The data were analyzed using FlowJo 10.4.2 (Tree Star, Ashland, OR) where viable cells were gated from a forward scatter/side scatter plot and singlet inclusion. Following neutrophil (Ly6G^hi^) and subsequent macrophage (CD11c^hi^F4/80^hi^) exclusion, CD8^+^ T cells were gated as CD3e^+^DX5^−^CD4^−^CD8*α*^+^.

To distinguish blood borne CD8^+^ T cells from those in the lung parenchyma, we used antibodies CD3e (BV786, clone 145-2C11, Biolegend), CD4 (FITC, clone RM4-5, Biolegend), CD8*α* (BV605, clone 53-6.7, BD Biosciences), B220 (APCe780, clone RA3-6B2, eBioscience), CD49b (APCe780, clone DX5, eBioscience), CD62L (PE-Cy7, clone MEL-14, Biolegend), CD69 (PE, clone H1.2F3, Biolegend), CD44 (PE-Dazzle594, clone IM7, Biolegend), CD25 (Alexa700, clone PC61, Biolegend), CD43 (APC, clone 1B11, Biolegend), CD127 (PE-Cy5, clone A7R34, Biolegend), and CD103 (BV711, clone 2E7, Biolegend) or CD314 (NKG2D) (BV711, clone CX5, BD Bioscience). The data were analyzed using FlowJo 10.6.2 (Tree Star, Ashland, OR) where viable cells were gated from a forward scatter/side scatter plot, singlet inclusion, and viability dye exclusion. Blood borne CD8^+^ T cells were gated as CD45^+^ cells and those in the lung parenchyma as CD45^−^. The total in sub-population were gated as CD3e^+^B220^−^DX5^−^CD4^−^CD8^+^. IAV-specific CD8^+^ T cells were then gated as CD25^+^CD43^+^ (recently activated), CD25^−^CD43^+^ (effector) [46], and CD43^−^CD127^+^CD103^+^CD69^+^ (long-lived memory). Expression of CD44, CD69, CD62L, and NKG2D were also assessed to ensure appropriate classification.

For all experiments, the absolute numbers of CD8^+^ T cells were calculated based on viable events analyzed by flow cytometry as related to the total number of viable cells per sample. We use the kinetics of the total number of CD8^+^ T cells (non-perfused lung; “total”), the total number of CD8^+^ T cells in the lung parenchyma (perfused lung; “lung-specific”), and specific virus-specific CD8^+^ T cell phenotypes in the lung parenchyma as defined by surface staining (described above). We chose this approach because the use of tetramers yields epitope-specific cell dynamics that vary in time and magnitude (e.g., as in [13, 46]) and would complicate the model and potentially skew the results. In addition, tetramer staining is not available for all epitopes in BALB/cJ mice and they may underestimate the dynamics of IAV-specific cells [46]. The gating strategies are shown in Supplementary file 1F.

### Lung immunohistopathologic and immunohistochemical (IHC) evaluation

The lungs from IAV infected mice were fixed via intratracheal infusion and then immersion in 10% buffered formalin solution. Tissues were paraffin embedded, sectioned, and stained for influenza virus using a primary goat polyclonal antibody (US Biological, Swampscott, MA) against influenza A, USSR (H1N1) at 1:1000 and a secondary biotinylated donkey anti-goat antibody (sc-2042; Santa Cruz Biotechnology, Santa Cruz, CA) at 1:200 on tissue sections subjected to antigen retrieval for 30 minutes at 98°C. The extent of virus spread was quantified by capturing digital images of whole-lung sections stained for viral antigen using an Aperio ScanScope XT Slide Scanner (Aperio Technologies, Vista, CA) then manually outlining defined fields. Alveolar areas containing virus antigen-positive pneumocytes were highlighted in red (defined as “active” infection), whereas lesioned areas containing minimal or no virus antigen-positive debris were highlighted in green (defined as “inactive” infection). Lesions containing a mix of virus antigen-positive and antigen-negative pneumocytes were highlighted in orange (defined as “mixed” infection). The percentage of each defined lung field was calculated using the Aperio ImageScope software. Pulmonary lesions in HE-stained histologic sections were assigned scores based on their severity and extent. Representative images and quantitative analyses of viral spread and lung pathology during IAV infection are shown in Figure 4.

### CD8^+^ T cell model

To examine the contribution of CD8^+^ T cells to the biphasic viral load decay, we expanded the density-dependent (DD) model (Equations (A9)–(A12)) to include two mechanisms of infected cell clearance (non-specific clearance (*δ*) and CD8^+^ T cell-mediated clearance (*δ_E_*(*I*_2_,*E*))) and two CD8^+^ T cell populations: effector (*E*, denoted CD8_E_) and memory (*E_M_*, denoted CD8_M_) CD8^+^ T cells. The model is given by Equations (1)–(6).

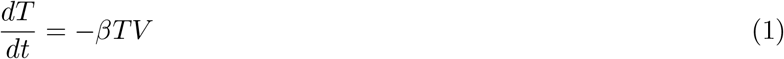

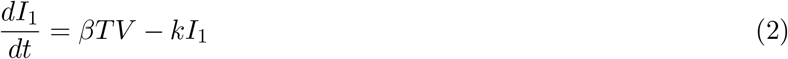

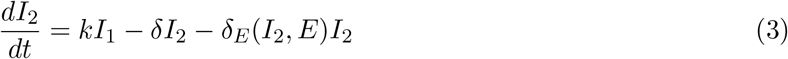

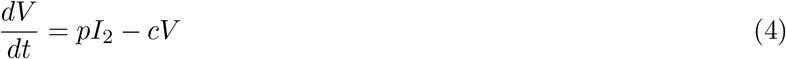

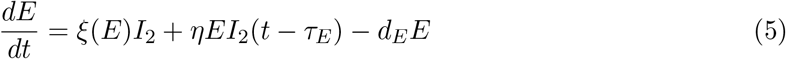

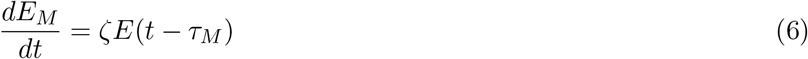

In this model, target cells become infected with virus at rate *βV* per day. Once infected, cells enter an eclipse phase (*I*_1_) before transitioning at rate *k* per day to a productively-infected state (*I*_2_). These infected cells produce virus at rate *p* TCID_50_ per infected cell per day, and virus is cleared at rate c per day. Virus-producing infected cells (*I*_2_) are cleared by non-specific mechanisms (e.g., apoptosis and/or innate immune responses) at a constant rate *δ* per day. Innate immune responses were excluded from the model because the viral load data is linear during the time where they act (2–7 d pi) and, thus, additional equations cannot improve the fit. Productively infected cells are cleared by CD8_E_ at rate *δ_E_*(*I*_2_,*E*) = *δ_E_E*/(*K_δ_E__* + *I*_2_) per day, where the rate of infected cell clearance is *δ_E_*/*K_δ_E__* per CD8_E_ per day and *K_δ_E__* is the half-saturation constant. The CD8_E_-mediated clearance rate (*δ_E_*(*I*_2_, *E*)) is dependent on the density of infected cells and is similar to the infected cell clearance term in the DD model (see Equation (A11)) [15]. Similar density-dependent forms have also been used in models that describe the CD8^+^ T cell response to other virus infections [33, 35, 98]. Models that exclude this density-dependence were examined, but these models resulted in a statistically poor fit to the data as defined by the Akaike Information Criteria (AIC) (Figure A1, Supplementary file 1G). This is due in part to the increase of CD8^+^ T cells from 3–5 d pi (Figure 1). We did examine a model that excluded these dynamics and included CD8^+^ T cell expansion as a density-dependent function (e.g., *ηEI*_2_(*t* – *τ_E_*)/(*K_I_* +*I*_2_)) while keeping a linear rate of CD8-mediated infected cell clearance (*δ_E_EI*_2_) (Equations (A1)–(A6); Figure A1). This model was adapted from other published models [40–42] and can produce similar dynamics from 6-10 d pi, but it was not statistically supported by our data (Supplementary file 1G). We further tested the model in [40], but this model could not fit our data and lacked statistical support (Figure A1, Supplementary file 1G).

The model assumes that the initial CD8_E_ influx in the lung is proportional to infected cells at rate *ξ*(*E*) = *ξ*/(*K_E_* + *E*) CD8_E_ per cell per day, which is down-regulated by the CD8_E_ already present in the lung. The associated half-saturation constant is *K_E_*. Similar terms for CD8_E_ regulation have been used in modeling HIV infections [35, 131] and in models that examined CD8^+^ T cell proliferation mechanisms [132]. We also examined whether their influx is proportional to infected cells at a delay (i.e., *ξ*(*E*)*I*(*t* – *τ_I_*)). While this modification better captured the initial increase in CD8^+^ T cells, the additional parameter was not supported. In our model, CD8_E_ expansion occurs at rate *η* per infected cell per day with time delay *τ_E_*. This term accounts for local CD8_E_ proliferation in the lung [45,133] and migration of CD8_E_ from secondary lymphoid organs [20,134–136]. The delay may signify the time it takes CD8_E_ to become activated by antigen presenting cells, differentiate, proliferate, and/or migrate to the infection site. The lung CD8_E_ population declines due to cell death and/or emigration at rate *d_E_* per day. These cells transition to CD8_M_ (*E_M_*) at rate *ζ* CD8_M_ per CD8_E_ per day after *τ_M_* days. The model schematic and fit to the viral load, total, effector, and memory CD8^+^ T cell data are in Figure 1.

### Inflammation model

To estimate the alveolar and interstitial inflammation without modeling other cell classes (e.g., macrophages and neutrophils), we assumed that inflammation in the lung (*L_I_*) was proportional to the infected cells (*I*_1_ and *I*_2_) according to the equation,

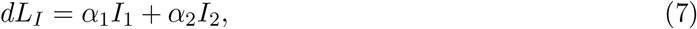

where *α*_1,2_ has units of score/cell/d and defines the inflammation score contribution from each infected cell class. A decay term was excluded because inflammation does not resolve on the timescale of our data.

### Parameter estimation

Given a parameter set *θ*, the cost *C*(*θ*) was minimized across parameter ranges using an Adaptive Simulated Annealing (ASA) global optimization algorithm [15] to compare experimental and predicted values of virus (*V*; log_10_ TCID_50_/lung) and log_10_ total CD8^+^ T cells/lung (*Ê* = *E*+*Ê_M_* + *Ê*_0_, where *Ê*_0_ is the initial number of CD8^+^ T cells at 0 d pi), or log_10_ TCID_50_/lung virus (*V*), log_10_ effector CD8^+^ T cells/lung (*Ê* = *E*+*Ê*_0_), and log_10_ memory CD8^+^ T cells/lung (*Ê_M_* = *E_M_*+*Ê*_*M*_0__). The cost function is defined by

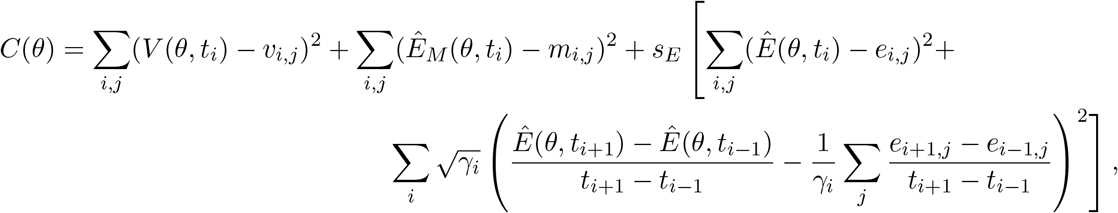

where (*t_i_, v_i,j_*) is the viral load data, (*t_i_, e_i,j_*) is the total or effector CD8^+^ T cell data, (*t_i_, m_i,j_*) is the memory CD8^+^ T cell data, and *V*(*θ,t_i_*;), *Ê*(*θ, t_i_*), and *Ê_M_*(*θ,t_i_*) are the corresponding model predictions. Here, *s_E_* = (*v*_max_ – *v_min_*)/(*e*_max_ – *e*_min_) is a scaling factor, and *γ_i_* = *J*_*i*+1_ *J*_*i*−1_ where *J_i_* is the number of observations at time *t_i_*. Errors of the log_10_ data were assumed to be normally distributed. To explore and visualize the regions of parameters consistent with the model, we fit Equations (1)–(6) to 2000 bootstrap replicates of the data. If the fit was within χ^2^ = 0.05 of the best-fit and the CD8 derivative was not a statistical outlier as determined by the function *isoutlier*, then the bootstrap was considered successful [15,128,137]. For each best-fit estimate, we provide 95% confidence intervals (CI) obtained from the bootstrap replicates (Table 1). Calculations were performed either in MATLAB using a custom built ASA algorithm [15] or in Python using the *simanneal* package [138] followed by a L-BFGS-B [139,140] deterministic minimization through SciPy’s *minimize* function. MATLAB *ode15s* and *dde23* or SciPy integrate.ode using *lsoda* and PyDDE [141] were used as the ODE and DDE solvers.

Estimated parameters in the CD8^+^ T cell model included the rates of virus infection (*β*), virus production (*p*), virus clearance (*c*), eclipse phase transition (*k*), non-specific infected cell clearance (*δ*), CD8_E_-mediated infected cell clearance (*δ_E_*), half-saturation constants (*K_δ_E__* and *K_E_*), CD8_E_ infiltration (*ξ*), CD8_E_ expansion (*η*), delay in CD8_E_ expansion (*τ_E_*), CD8_E_ clearance (*d_E_*), CD8_M_ generation (*ζ*), delay in CD8_M_ generation (*τ_M_*), and the baseline number of CD8^+^ T cells 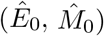. Bounds were placed on the parameters to constrain them to physically realistic values. Because biological estimates are not available for all parameters, ranges were set reasonably large based on preliminary results and previous estimates [15]. The rate of infection (*β*) was allowed to vary between 10^−6^ – 10^−1^ 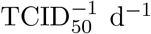, and the rate of virus production (p) between 10^−1^ – 10^3^ TCID_50_ cell^−1^ d^−1^. Bounds for the virus clearance rate (*c*) were 1 d^−1^ (*t*_1/2_ = 16.7 h) and 10^3^ d^−1^ (*t*_1/2_ = 1 min). To insure biological feasibility, the lower and upper bounds for the eclipse phase transition rate (*k*) were 4 – 6 d^−1^ as done previously [15].

The rate of non-specific infected cell clearance (*δ*) was given limits of 0.05 – 1 d^−1^. The CD8_E_-mediated infected cell clearance rate (*δ_E_*) varied between 0.01 – 2 cells 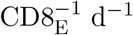, and the associated half-saturation constant (*K_δ_E__*) was bounded between 10^1^ – 10^6^ cells. The upper bound of *δ_E_* was chosen to maintain the convergence of *δ* to nonzero values. Bounds for the rate of CD8_E_ infiltration (*ξ*) were 10^2^ – 10^6^ 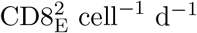, and bounds for the half-saturation constant (*K_E_*) were 10^3^ – 10^7^ CD8_E_. The CD8_E_ expansion rate (*η*) varied between 10^−8^ – 10^−6^ cell^−1^ d^−1^, and the delay in CD8_E_ expansion (*τ_E_*) between 2–6 d. The rate of CD8_E_ clearance (*d_E_*) had limits of 0.05 – 2 d^−1^. The rate of CD8_M_ generation (*ζ*) varied between 0.01 – 1 CD8_M_ CD8_E_^−1^ d^−1^, and the delay in CD8_M_ generation (*τ_M_*) varied between 3–4 d (total CD8^+^ T cell fit) or 2–6 d pi (effector and memory CD8^+^ T cell fit). Larger bounds were examined for this parameter, however, the parameter was non-identifiable in the total CD8^+^ T cell fit and a small range was required for convergence. Bounds for the baseline number of CD8^+^ T cells (*Ê*_0_, *Ê*_*M*_o__) were set to the upper and lower values of the data at 0 d pi (3.0 × 10^5^ – 5.3 × 10^5^ CD8 (total), 2.5 × 10^2^ – 1.4 × 10^3^ CD8_E_ (effector), 3.4 × 10^2^ – 8.0 × 10^2^ CD8_M_ (memory)).

The initial number of target cells (*T*(0)) was set to 10^7^ cells [15,128,137]. The initial number of infected cells *I*_1_(0) was set to 75 cells to reflect an initial dose of 75 TCID_50_ [15]. We previously found that estimating *I*_1_(0), fixing *V*(0) = 75 TCID_50_, or estimating *V*(0) did not improve the fit and could not be statistically justified [15]. The initial number of productively infected cells (*I*_2_(0)), the initial free virus (*V*(0)), and the initial number of CD8_E_ (E(0)) and CD8_M_ (*E_M_*(0)) were set to 0. All other parameter estimations were done as described in the text.

### Linear regression

The function *polyfit* in MATLAB was used to perform linear regression on the percent active lesioned area, the percent inactive lesioned area, and the CD8^+^ T cells during the expansion phase (5–8 d pi) and the contraction phase (9–10 d pi). Linear fits are shown in Supplementary file 1C.

### Cumulative area under the curve

The function *cumtrapz* in MATLAB was used to estimate the cumulative area under the curve (CAUC) of the infected cells (*I*_2_) for the best-fit model solution.

## 3 Acknowledgments

This work was supported by NIH grant AI100946, AI125324, and AI139088 and ALSAC. We thank Alan Perelson for his helpful comments, and Robert Michael for technical assistance.

## A Appendix

### A.1 Alternate CD8^+^ T cell Models

We examined alternate formulations of the CD8^+^ T cell model to further investigate the density-dependence in the CD8^+^ T cell response. Rather than assuming that CD8_E_-mediated clearance of infected cells is dependent on their density, the model in Equations (A1)–(A6) assumes that the rate of CD8_E_ expansion is dependent on the density of infected cells. Similar models have been used to study CD8^+^ T cell responses during HIV infection [41,42] and other viral infections [40]. The differences between this alternate model and the CD8^+^ T cell model (Equations (1)–(6)) are in bold.

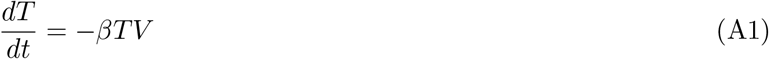

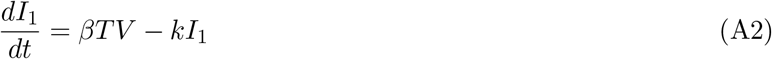

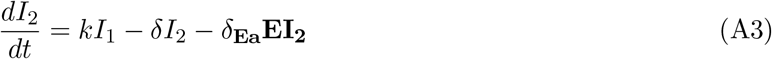

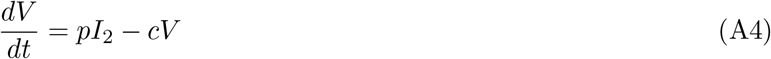

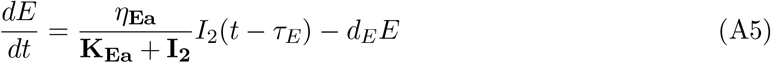

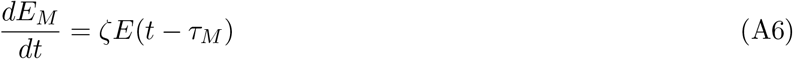

When a linear CD8_E_-mediated infected cell clearance rate is included, the CD8_E_ dynamics between 3–5 d pi cannot be replicated and returned a statistically worse fit via AIC (Supplementary file 1G). However, because these cells may not have effector functions and contribute to infected cell clearance at this time (see Discussion), we excluded these data and the term *ξI*_2_/(*K_E_* + *E*) when fitting Equations (A1)–(A6) to the viral load and total CD8^+^ T cell data (Figure A1). The CD8 dynamics are similar to those generated by CD8^+^ T cell model. However, the alternate model underestimates the data at day 7 and the sharp decline between 7–8 d. While it is inappropriate to directly compare these models due to the varying number of data points, we assessed their goodness of fit with and without contributions from the data at 3–5 d pi (Supplementary file 1G). Regardless of the data inclusion or exclusion, the alternate model had a higher AIC in all contexts and, thus, is not statistically justifiable. Similarly, we examined the model in [40] (Equations (A7)–(A8)), which also could not replicate our data (Figure A1).

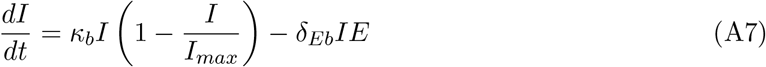

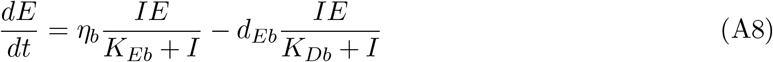

**Figure A1.**
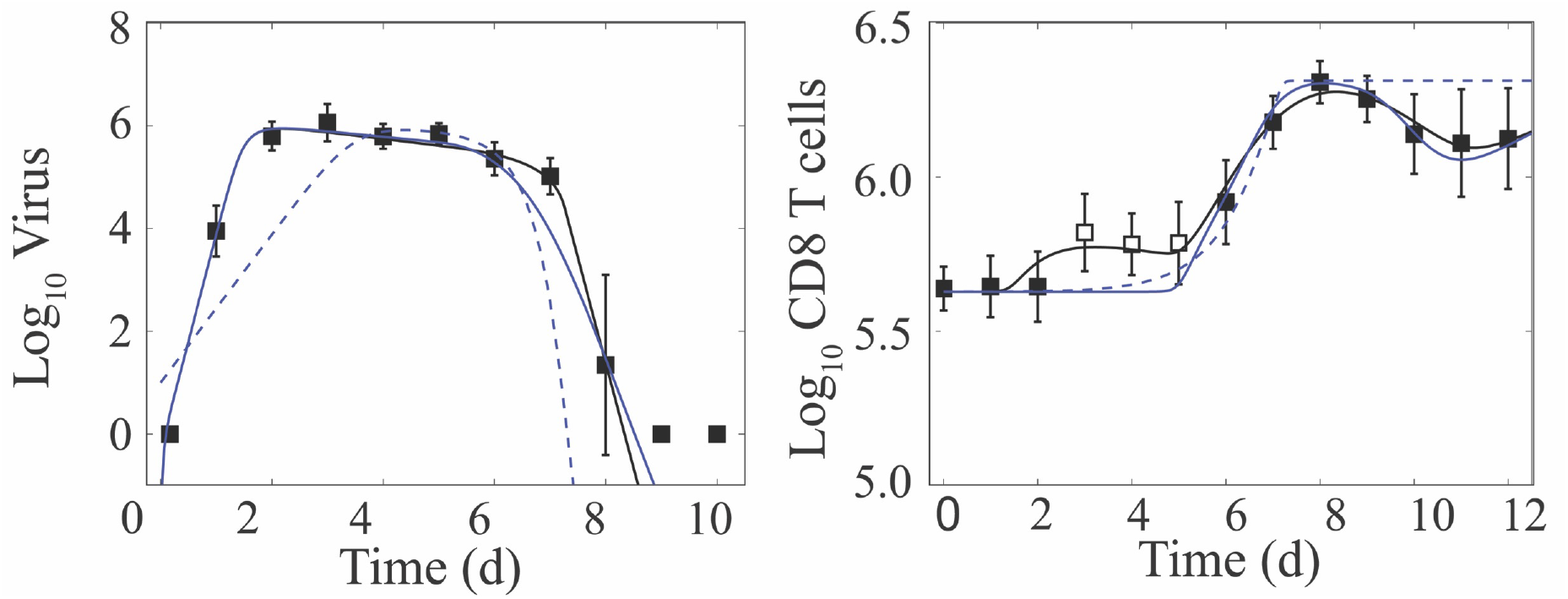
Fit of Alternate CD8^+^ T cell Models. Fit of the CD8^+^ T cell model (Equations (1)–(6); solid black line)) compared to the fit of two alternate CD8^+^ T cell models (Equations (A1)–(A6) (solid blue line) and Equations (A7)–(A8) (dashed blue line)) to virus and CD8^+^ T cells (excluding 3–5 d pi; white squares) from the lungs of mice infected with 75 TCID_50_ PR8 (10 mice per time point). Resulting parameter values were *δ_E_a__* = 4.02 × 10^−6^ CD8^−1^ d^−1^, *ηE_a_* = 3.12 × 10^5^ CD8_E_/d, and *K_Ea_* = 9.53 × 10^5^ infected cells, and *κ_b_* = 3.33 d^−1^, *I_max_* = 1.0 × 10^6^ infected cells, *δ_Eb_* = 1.53 × 10^−5^ CD8_E_^−1^ d^−1^, *η_b_* = 1.49 d^−1^, *K_Eb_* = 9.5 infected cells, *d_Eb_* = 0.95 d^−1^, and *K_Db_* = 1.90 × 10^6^ infected cells. All other parameters are in Table 1, and the AICs are in Supplementary file 1G. Data are shown as mean ± standard deviation.

### A.2 Comparison of the Density-Dependent and CD8^+^ T cell Models

We previously developed a density-dependent (DD) viral kinetic model, which describes the biphasic decline of viral loads without inclusion of specific host responses [15]. This model tracks 4 populations: susceptible epithelial (“target”) cells (*T*), two classes of infected cells (*I*_1_ and *I*_2_), and virus (*V*) [15].

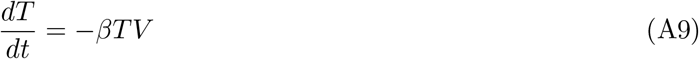

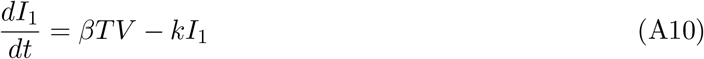

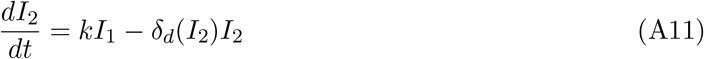

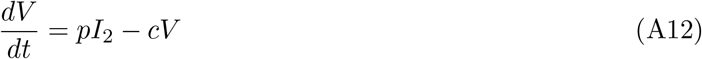

Briefly, in the DD model, virus-producing infected cells (*I*_2_) are cleared according to the function *δ_d_*(*I*_2_) = *δ_d_*/(*K_δ_* + *I*_2_), where *δ_d_*/*K_δ_* is the maximum per day rate of infected cell clearance and *K_δ_* is the half-saturation constant (Figure A2). All other terms are common to the CD8 T^+^ cell model (Equations (1)–(6)). The DD model provides a close fit to the viral load data in Figure 1B and replicates the biphasic viral load decline while excluding the dynamics of specific immune responses [15]. Unsurprisingly, the CD8^+^ T cell model is also capable of reproducing the biphasic viral load decay (Figure 1B and A2). In that model, infected cell clearance is split into terms for non-specific clearance (*δ*) and CD8_E_-mediated clearance (*δ_E_*(*I*_2_,*E*) = *δ_E_E*/(*K_δ_* + *I*_2_)) (Figure A2).

Because the CD8^+^ T cell model is more mechanistic than the DD model, most of the correlations between the parameters common to both models (i.e., the rates of virus infectivity (*β*), virus production (*p*), and virus clearance (*c*)) were reduced (Figure A2A). In addition, the correlations between the infected cell clearance parameters (*δ_d_* and *K_δ_* or *δ_E_* and *K_δ_E__*) and between the rate of virus infectivity (*β*) and their ratios (*δ_d_*/*K_δ_* or *δ_E_*/*K_δ_E__*) were abolished (Supplementary file 1B). There was a negative correlation between the infected cell clearance parameters (*δ* and *δ_E_*; Figure 2B), which may reflect the connection between the efficacy of early immune mechanisms and the CD8^+^ T cell response. This result is in line with experimental evidence that the innate immune responses modulate the activation of adaptive immunity [142–146].

The differences in model structure between the two models yielded changes in parameter sensitivity and model behavior during the rapid viral clearance phase (Figure A2). In the DD model, the most sensitive parameter is the infected cell clearance, *δ_d_* (Figure A2). A 50% decrease in this parameter resulted in a ~7 d delay in viral resolution (Figure A2) [15]. In the CD8^+^ T cell model, however, viral resolution is delayed by <1 d if the CD8_E_-mediated infected cell clearance parameter (*δ_E_*) is reduced by 50% (Figures A2–A3). The rates of CD8_E_ expansion (*η*) and decay (*d_E_*) are sensitive and, thus, significantly influence the viral resolution kinetics (Figures A2–A4). A 50% decrease in *η* results in a ~6 d delay in recovery (Figures A2–A3) whereas a 48% decrease in *η* prolongs the infection by ~30 d (Figure 3D–E). This bifurcation in recovery time is a unique feature of the CD8^+^ T cell model.

**Figure A2.**
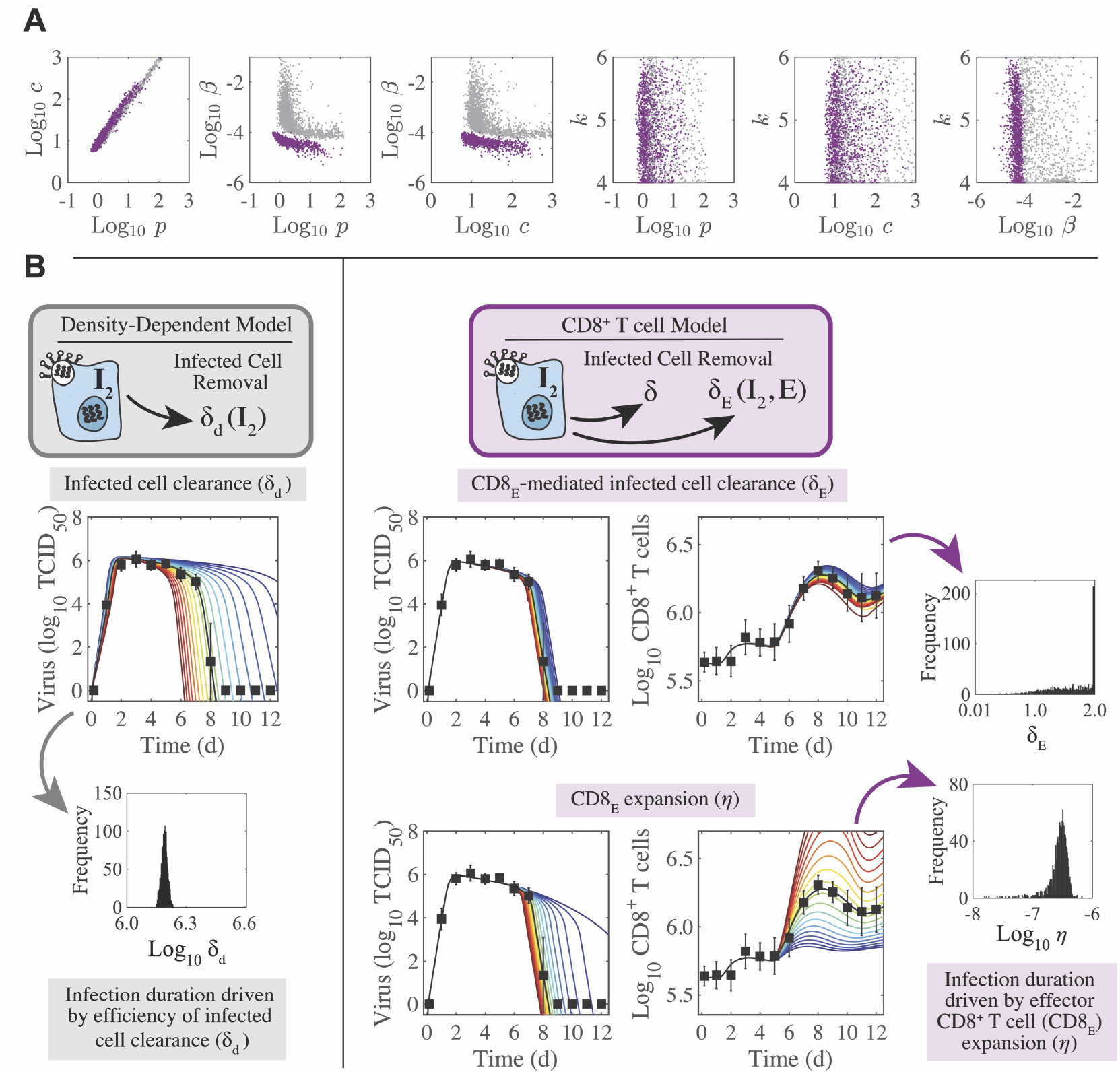
Parameter behavior of the density-dependent model and the CD8^+^ T cell model. (A) Comparison of parameters that were common between the density-dependent model (gray, Equations (A9)–(A12)) and the CD8^+^ T cell model (purple, Equations (1)–(6)). Correlations were evident between parameters relating to the rates of virus infectivity (*β*), virus production (*p*), and virus clearance (*c*). However, the strength of the correlation was significantly reduced in the CD8^+^ T cell model. The eclipse phase parameter (*k*) was not well-defined in either model. (B) In the density-dependent model (gray), the viral kinetics and the infection duration were sensitive to small changes in the infected cell clearance parameter (*δ_d_*). This parameter was well-defined with a narrow 95% CI. In the CD8^+^ T cell model (purple), changing the CD8_E_-mediated infected cell clearance parameter (*δ_E_*) had little impact on viral kinetics or CD8^+^ T cell kinetics. However, these kinetics were most sensitive to changes in the rate of CD8_E_ expansion (*η*), which was well-defined with a narrow 95% CI.

### A.3 Regulation of the CD8^+^ T cell Response

To further understand the regulation of the CD8^+^ T cell response, we examined the 2-D parameter ensembles (Figure 2A–C, Supplementary file 1B) and the results from the sensitivity analysis (Figure A3–A4). Overall, few parameters were correlated. There was an expected, although small, positive correlation between the rate of CD8_E_ infiltration (*ξ*) and the associated half-saturation constant (*K_E_*) (Supplementary file 1B), which represents the coordination between CD8_E_ recruitment and the processes that prevent an overabundance of these cells. Likewise, a negative correlation was detected between the rate of initial CD8_E_ influx (*ξ*) and the initial number of CD8^+^ T cells (*Ê*_0_) (Supplementary file 1B). The influx rate (*ξ*) was also positively correlated with the delay in CD8_E_ expansion (*τ_E_*) (Supplementary file 1B). The rates of CD8_E_ expansion (*η*) and decay (*d_E_*) are correlated (Figure 2C), indicating a balance between these two processes. This correlation was expected and reflects the coordination of mechanisms that regulate CD8^+^ T cell numbers, which may be necessary to limit excessive immunopathology while still resolving the infection [5, 6, 9]. Further, because of this correlation and the sensitivity of *η* (Figure A3), the CD8^+^ T cell kinetics are sensitive to changes in *d_E_* (Figure A4). However, increasing the decay rate had less impact on the viral load kinetics, comparatively. Because *d_E_* is correlated with both *η* and the rate of CD8_M_ generation (*ζ*) (Figure 2C), it naturally follows that *η* and *ζ* are correlated (Supplementary file 1B). Changing the rates of virus infectivity (*β*), production (*p*), or clearance (*c*) had little effect on viral load or CD8^+^ T cell kinetics (Figure A3).

**Figure A3.**
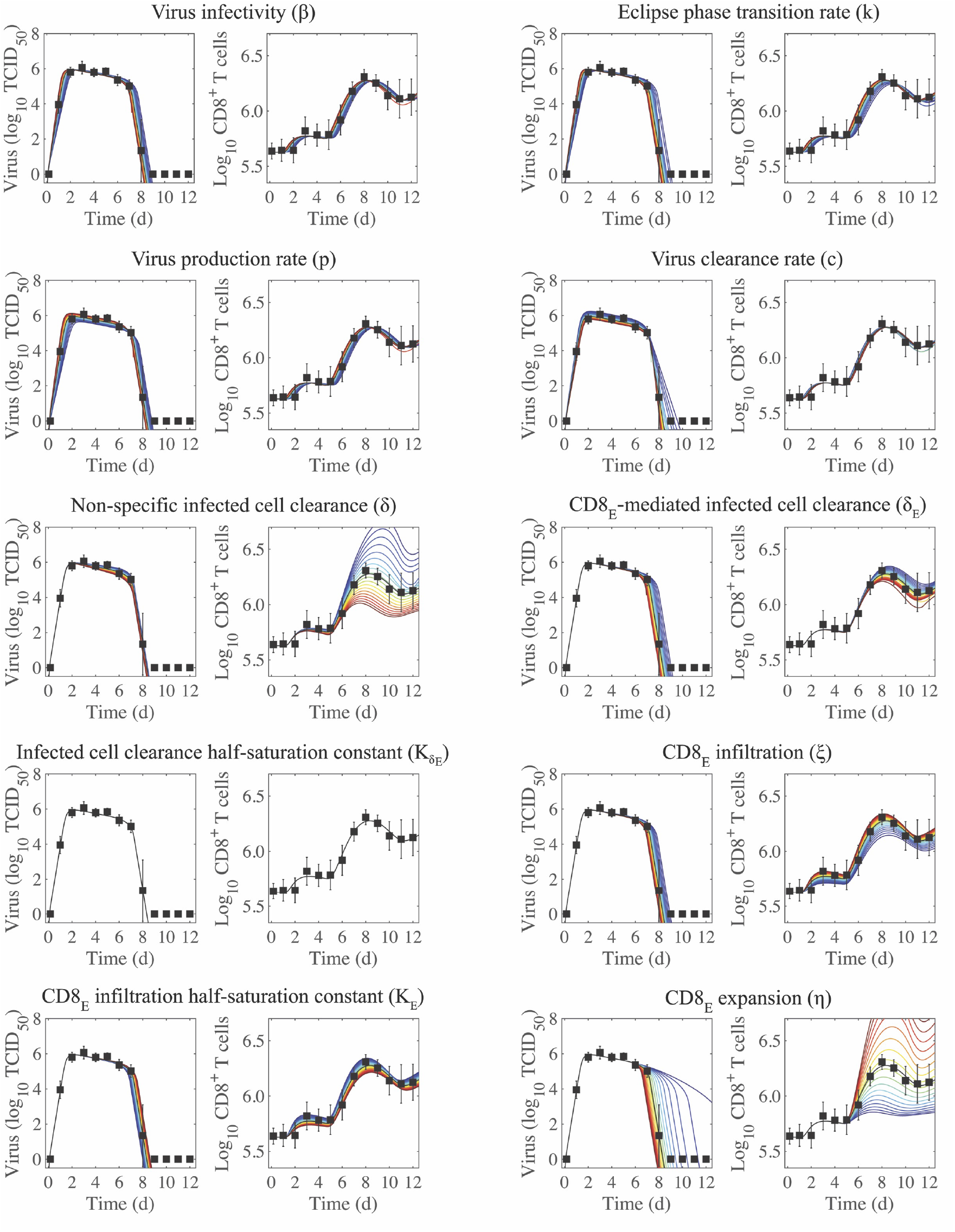
Sensitivity of the CD8^+^ T cell model. Solutions of the CD8^+^ T cell model (Equations (1)–(6)) with the indicated parameter (*β, k, p, c, δ, δ_E_, *K_E_*, ξ, K_E_*, or *η*) increased (red) or decreased (blue) 50% from the best-fit value (Table 1). CD8_E_ denotes effector CD8^+^ T cells.

**Figure A4.**
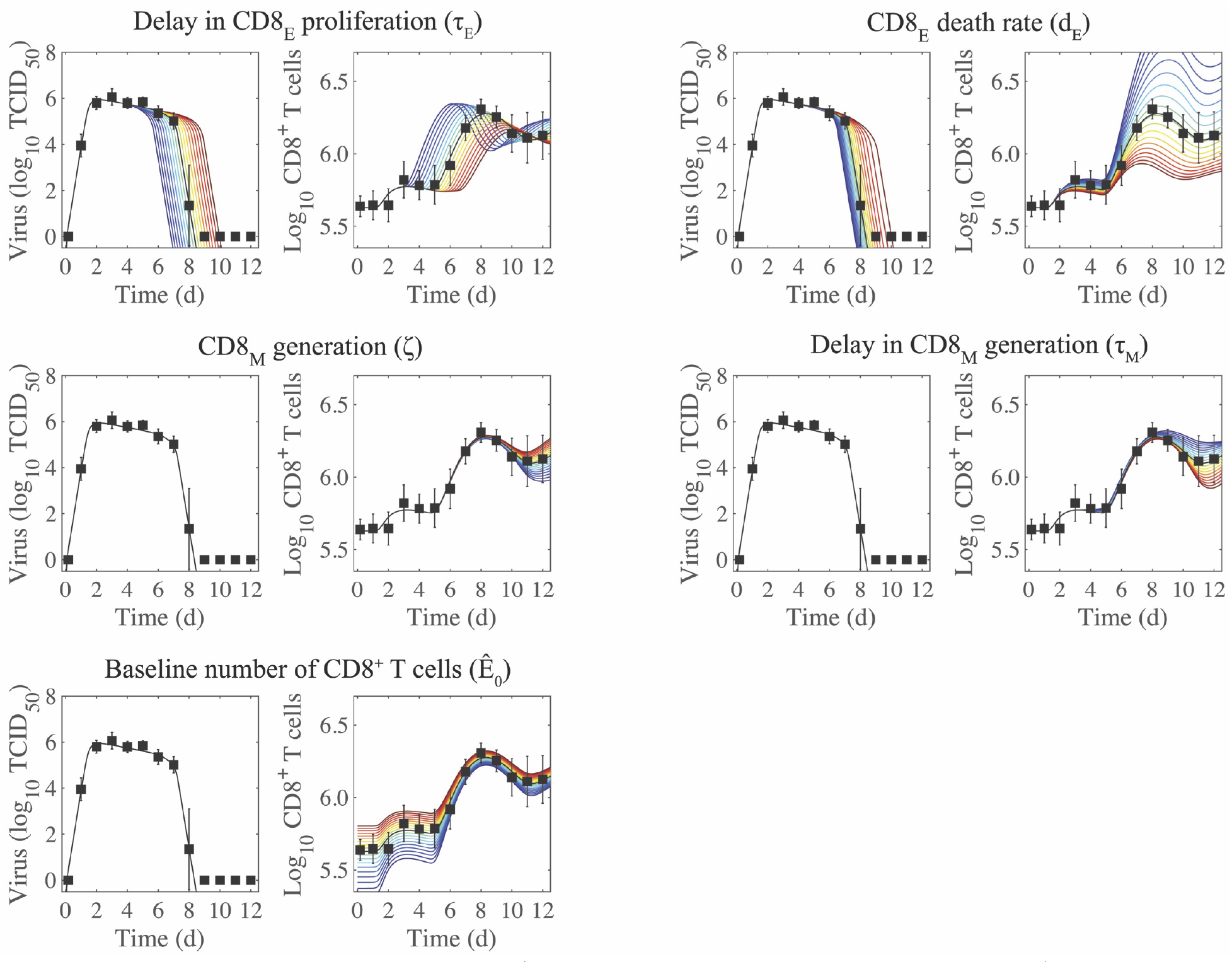
Sensitivity of the CD8^+^ T cell model. Solutions of the CD8^+^ T cell model (Equations (1)–(6)) with the indicated parameter (*τ_E_, d_E_, ζ, τ_M_*, or *Ê*_0_) increased (red) or decreased (blue) 50% from the best-fit value (Table 1). CD8_E_ and CD8_M_ denote effector and memory CD8^+^ T cells, respectively.

## Notes

### Competing Interest Statement

The authors have declared no competing interest.

### Summary of Updates

Substantial new data, analyses, and text have been included to validate the model, data, and method.

