## Supplementary File for "Dynamically Linking Influenza Virus Infection Kinetics, Lung Injury, Inflammation, and Disease Severity"

#### Supplementary file 1A

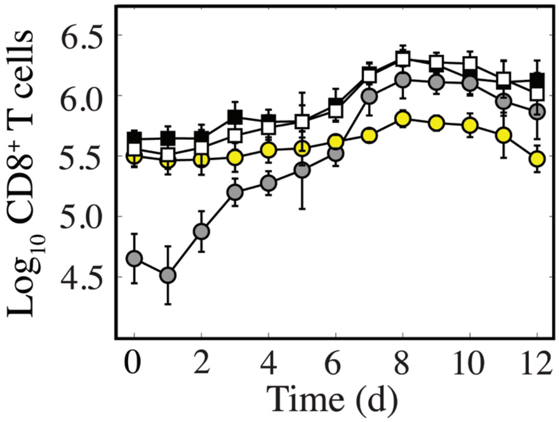


**Figure S1. Dynamics of CD8$^+$ T cells in the lung parenchyma and vasculature.** Comparison of total CD8$^+$ T cells in a non-perfused lung (black) and in a perfused lung (white). Of those from a perfused lung, comparison of CD8$^+$ T cells located in the lung parenchyma (CD45$^-$; gray) and vasculature (CD45$^+$; yellow). These data show that perfusion does not alter the number of cells recovered, and that the total number of cells primarily reflects changes in the lung parenchyma with little fluctuation in the lung vasculature. The cells in the lung vasculature may be slightly overestimated late in the infection due to incomplete perfusions during the height CD8-mediated clearance (7-9 d pi) when the lung integrity is low.

#### Supplementary file 1B

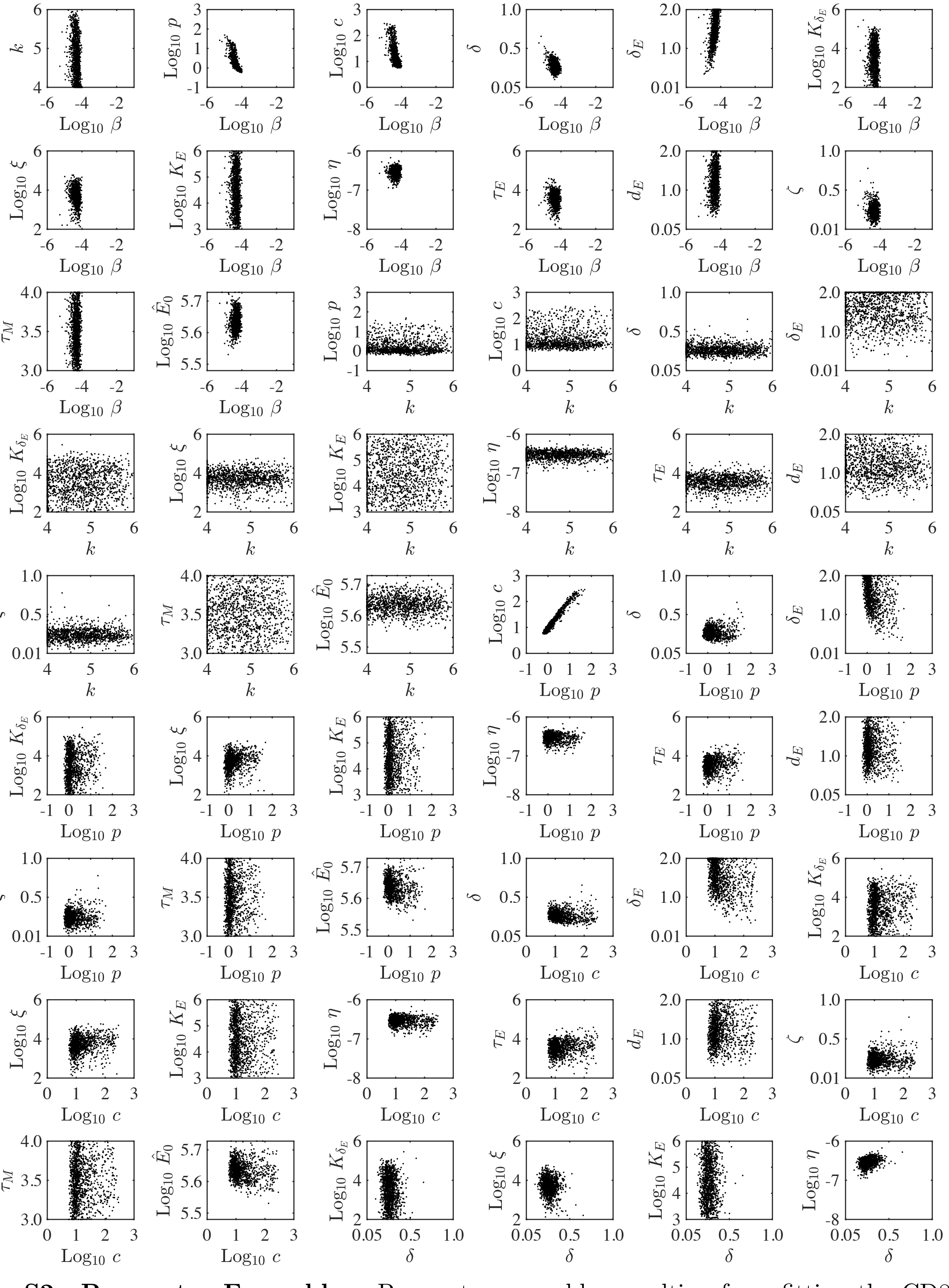


**Figure S2. Parameter Ensembles.** Parameter ensembles resulting from fitting the $CD8^+$ T cell model (Equations (1)–(6), Main Text) to viral loads and $CD8^+$ T cells from mice infected with 75 $TCID_{50}$ PR8. The axes limits reflect the imposed bounds. Additional ensemble plots are in Figure 2 (Main Text) and Figure S3.

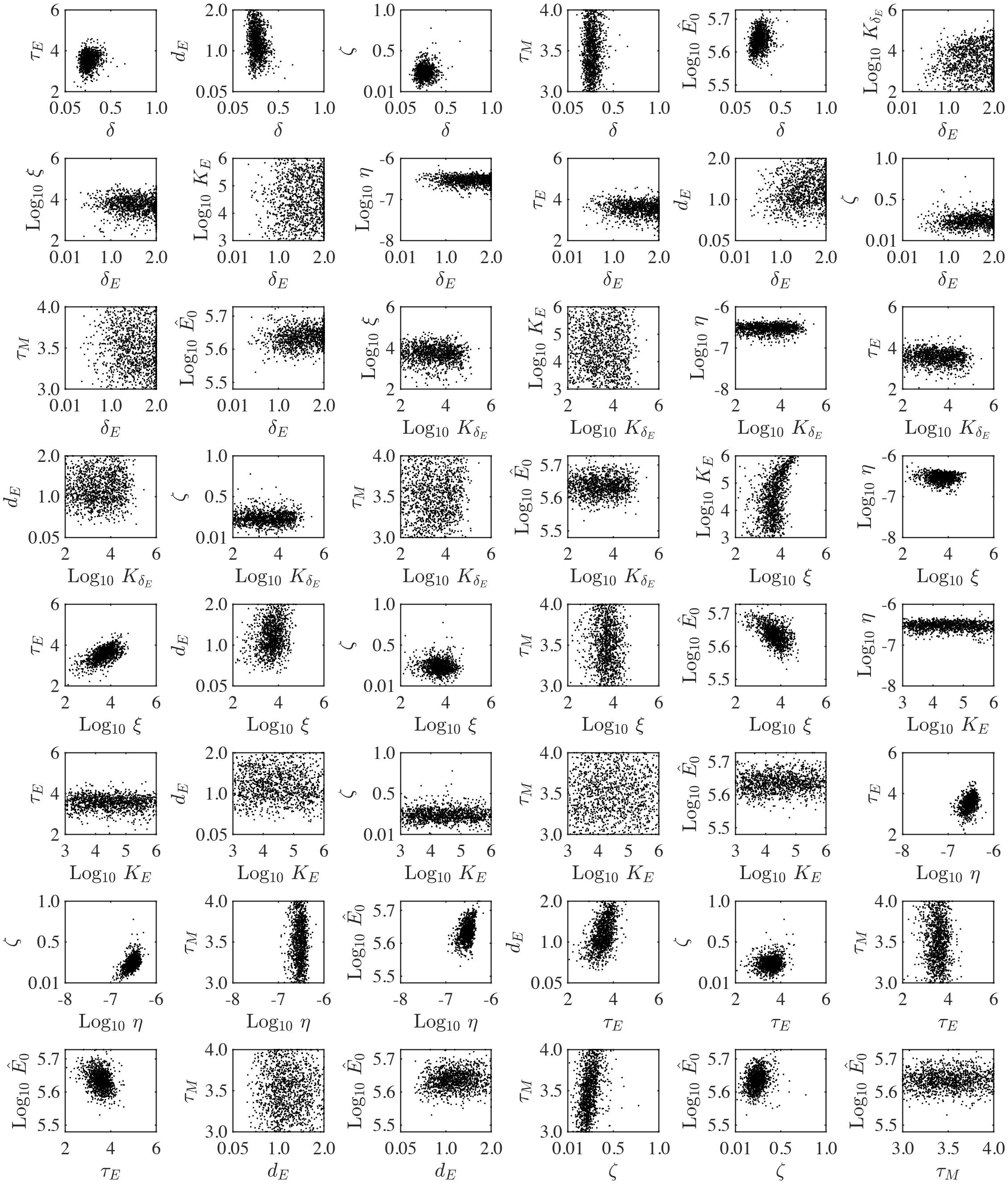


**Figure S3. Parameter Ensembles.** Parameter ensembles resulting from fitting the $CD8^+$ T cell model (Equations (1)–(6), Main Text) to viral loads and $CD8^+$ T cells from mice infected with 75 $TCID_{50}$ PR8. The axes limits reflect the imposed bounds. Additional ensemble plots are in Figure 2 (Main Text) and Figure S2.

#### Supplementary file 1C

**Table S1. $CD8^+$ T cell depletion model parameters.** Parameters, SSR, and $AIC_c$ obtained from fitting the $CD8^+$ T cell model (Equations (1)–(6)) to viral titers and $CD8^+$ T cells from mice infected with 75 $TCID_{50}$ PR8 and with CD8s depleted at -2, 0, 3, and 7 d pi. The total number of $CD8^+$ T cells is $\hat{E} = E + E_M + \hat{E}_0$, where $\hat{E}_0 = 5.6e3$ cells, and all other parameters are those in Table 1. The best model is bolded.

| $T_0$ | $p$ | $\delta$ | $c$ | $\beta$ | $\delta_E$ | $\xi$ | $\eta$ | $d_E$ | $K_\delta$ | $\tau_E$ | **SSR** | **$AIC_c$** |
|---|---|---|---|---|---|---|---|---|---|---|---|---|
| 3.17e6 | | | | | | | | | | | 342.34 | 60.86 |
| | 0.39 | | | | | | | | | | 320.36 | 60.19 |
| | | 1.09 | | | | | | | | | 177.95 | 54.31 |
| | | | 34.95 | | | | | | | | 295.88 | 59.40 |
| | | | | 1.29e-4 | | | | | | | 62.93 | 43.92 |
| | | | | | 2.63 | | | | | | 52.72 | 42.15 |
| | | | | | | 3.81e4 | | | | | 63.13 | 43.95 |
| | | | | | | | 2.81e-7 | | | | 52.24 | 42.06 |
| | | | | | | | | 0.84 | | | 53.51 | 42.30 |
| | | | | | | | | | 3.26e-8 | | 346.68 | 60.98 |
| | | | | | | | | | | 3.20 | 60.74 | 43.57 |
| 3.57e6 | | | | | | | 9.91e-7 | | | | 5.39 | 22.57 |
| | | | | 2.24e-5 | 6.04 | | | | | | 18.40 | 34.84 |
| | | 0.45 | | | | | 3.58e-7 | | | | 20.11 | 35.72 |
| | | 0.42 | | | 5.19 | | | | | | 20.53 | 35.93 |
| | | | | | | 4.29e3 | 4.14e-7 | | | | 25.72 | 38.19 |
| 7.97e6 | | | | | 4.48 | | | | | | 48.50 | 44.53 |
| | | 0.22 | | | 2.63 | | | | | | 52.54 | 45.33 |
| 1.00e7 | | | | | | 3.28e4 | | | | | 61.30 | 46.87 |
| 3.98e6 | | | | | | 1.33e4 | 1.01e-6 | | | | 2.44 | 18.92 |
| | | | 27.88 | | | 6.20e3 | 4.21e-7 | | | | 7.33 | 29.92 |
| | 0.51 | | | | | 1.04e4 | 4.15e-7 | | | | 15.22 | 37.23 |
| | 0.40 | 0.14 | | | 5.55 | | | | | | 16.46 | 38.01 |
| | | 0.34 | | | | 1.85e3 | 5.00e-7 | | | | 17.39 | 38.56 |
| | | | | 2.21e-05 | 7.98 | 1.95e4 | | | | | 17.45 | 38.59 |
| | 0.47 | 0.28 | | | | | 3.55e-7 | | | | 18.20 | 39.02 |
| | | 0.23 | | 2.24e-5 | 6.08 | | | | | | 18.38 | 39.11 |
| | 0.44 | 0.26 | | | | 8.10e4 | | | | | 44.31 | 47.91 |
| | 0.48 | 0.37 | | | | 4.57e3 | 5.18e-7 | | | | 2.40 | 24.75 |
| | | 0.44 | | 2.54e-5 | | 5.92e3 | 5.44e-7 | | | | 4.35 | 30.70 |
| 3.62e6 | | 0.24 | | | 1.63 | | 1.01e-6 | | | | 4.80 | 31.69 |

### Supplementary file 1D

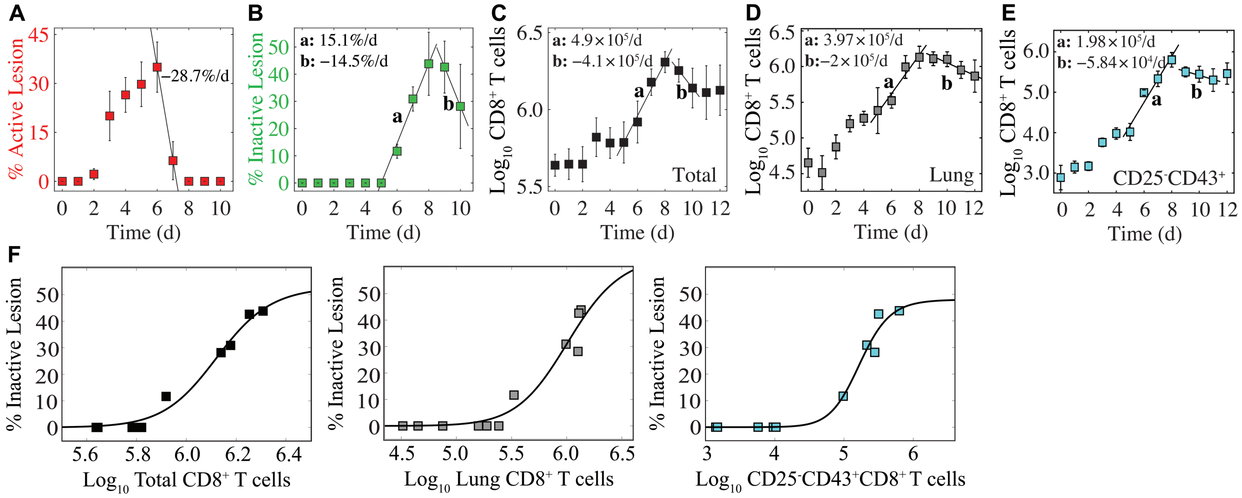


**Figure S4. Regression analysis of lung injury dynamics and $CD8^+$ T cells.** (A) Percent active lesion area decreases by 28.7%/d from 6–7 d pi. (B) Percent inactive lesion area increases by 15.1%/d from 5–8 d pi, and decreases by 14.5%/d from 9–10 d pi. (C) Total $CD8^+$ T cells increase at a rate of $4.9 \times 10^5$ cells/d from 5–8 d pi, and decrease at a rate of $4.1 \times 10^5$ cells/d from 9–10 d pi. (D) Lung $CD8^+$ T cells increase at a rate of $3.97 \times 10^5$ cells/d from 5–8 d pi, and decrease at a rate of $2 \times 10^5$ cells/d from 9–12 d pi. (E) $CD25^-CD43^+CD8^+$ T cells increase at a rate of $1.98 \times 10^5$ cells/d from 5–8 d pi, and decrease at a rate of $5.84 \times 10^4$ cells/d from 9–11 d pi. (F) Fit of a Hill function ($L_I = c_{max}E^n/(K_L^n + E^n)$) to the total $CD8^+$ T cells (black; $c_{max}$=52.88, $K_L$=6.12, $n$=56.32), Lung $CD8^+$ T cells (gray; $c_{max}$=64.65, $K_L$=6.02, $n$=25.16), and $CD25^-CD43^+CD8^+$ T cells (cyan; $c_{max}$=48.04, $K_L$=5.23, $n$=24.03).

#### Supplementary file 1E

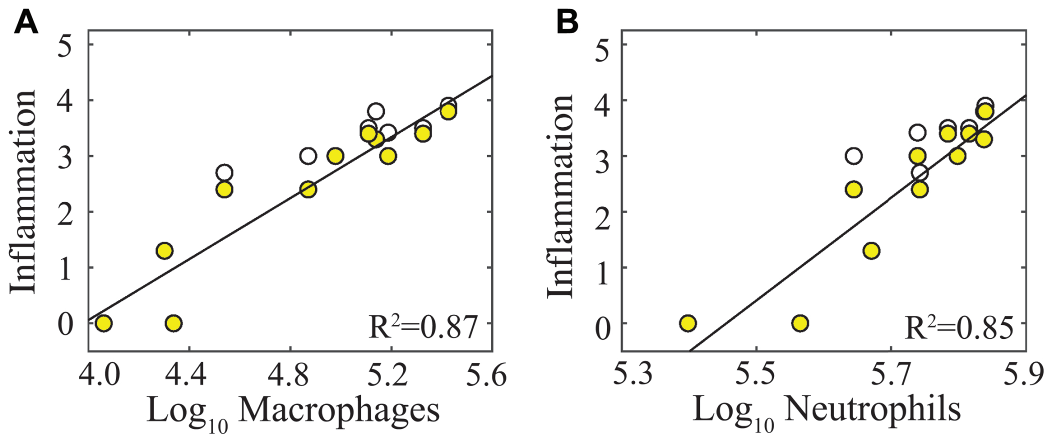


**Figure S5. Correlation of inflammation with macrophages and neutrophils.** Regression analysis of alveolar inflammation (white) and interstitial inflammation (yellow) with (A) $\log_{10}$ inflammatory macrophages ($F480^{hi}CD11c^{hi}CD11b^{+}$) and (B) $\log_{10}$ neutrophils ($Ly6G^{hi}$).

#### Supplementary file 1F

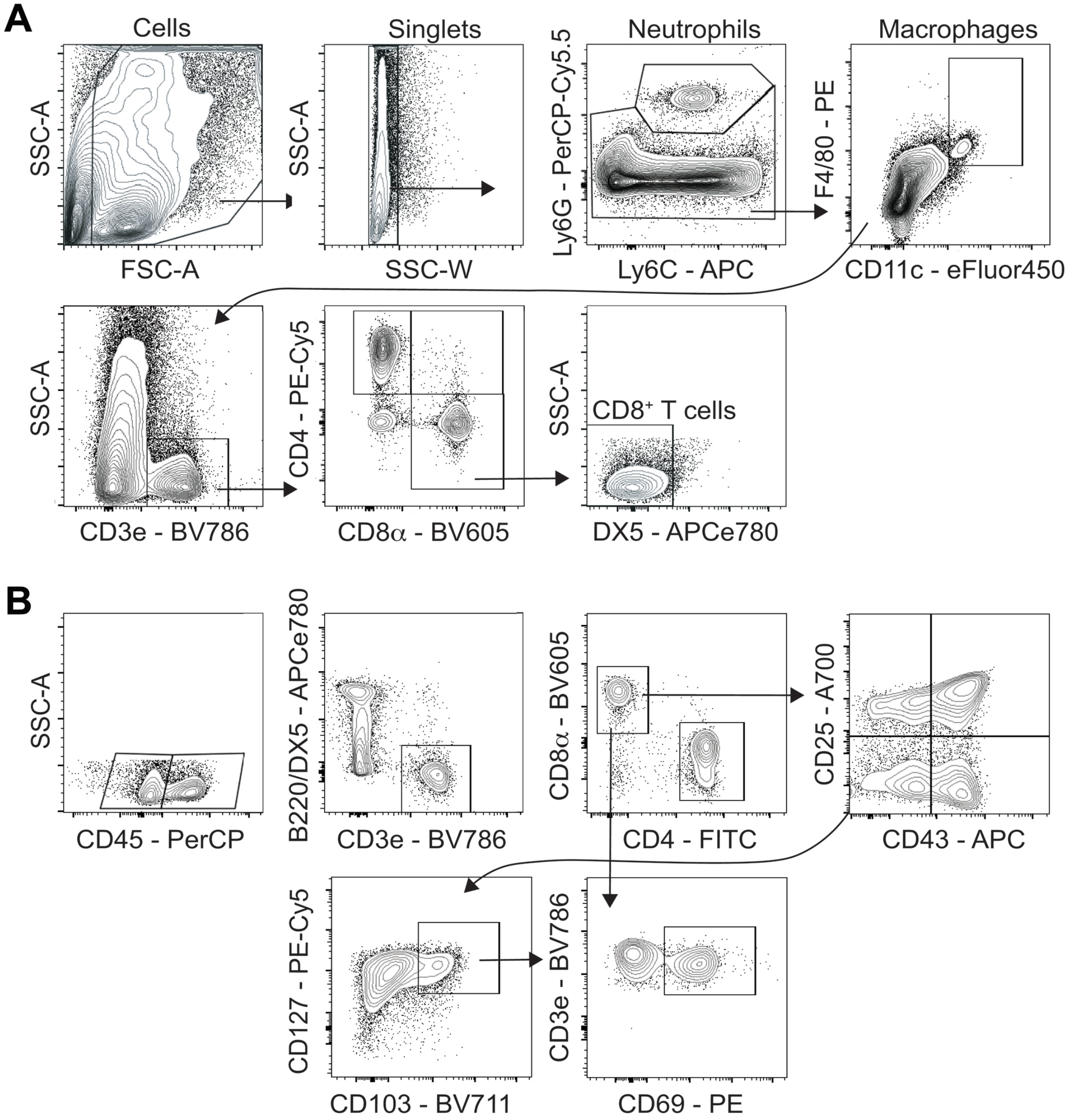


**Figure S6. Flow cytometry gating strategy for CD8$^+$ T cell analysis.** Live cells were first gated on forward scatter (FSC-A) and side scatter (SSC-A) then as singlets. (A) Following neutrophil (Ly6G$^{hi}$) and macrophage (F4/80$^{hi}$CD11c$^{hi}$) exclusion, T cells were gated as CD3e$^+$ with total CD8$^+$ T cells sub-gated as DX5$^-$CD4$^-$CD8$\alpha^+$. (B) Blood borne CD8$^+$ T cells were gated as CD45$^+$ cells and those in the lung parenchyma as CD45$^-$ with the total in each sub-population gated as CD3e$^+$B220$^-$DX5$^-$CD4$^-$CD8$^+$. IAV-specific CD8$^+$ T cells were sub-gated as CD25$^+$CD43$^+$ (recently activated), CD25$^-$CD43$^+$ (effector), and CD25$^-$CD43$^-$CD127$^+$CD103$^+$CD69$^+$ (long-lived memory). Expression of CD44, CD69, CD62L, and NKG2D were also assessed to ensure appropriate classification.

### Supplementary file 1G

**Table S2. Statistical comparison of alternate models.** Comparison of the Akaike Information Criteria (AIC) of the $CD8^+$ T cell model in Equations (1)–(6), the alternate model Equations (A1)–(A6), and the Baral model in Equations (A7)–(A8). The fit of these models is shown in Figure A1.

| | **Total CD8** | | **Lung CD8** | |
|---|---|---|---|---|
| | **All data** | **Excl. d3-5** | **All data** | **Excl. d3-5** |
| CD8 Model (Eqns (1)–(6)) | 32.1 | 26.0 | 77.9 | 63.2 |
| Alternate Model (Eqns (A1)–(A6)) | 69.5 | 64.9 | 84.7 | 72.5 |
| Baral Model (Eqns (A7)–(A8)) | 151.9 | 135.2 | 118.2 | 103.2 |
